# Ex vivo activation unmasks a sex-convergent, exhaustion-associated CD8+ T cell expansion in Parkinson’s disease

**DOI:** 10.64898/2026.08.13.744357

**Authors:** Friederike Grandke, Caroline Diener, Bianca Becher, Anouck Becker-Dorison, Nicole Ludwig, Barbara Walch-Rückheim, Tanja Tänzer, Klaus-Ulrich Dillmann, Martin Hart, Marcus Unger, Klaus Faßbender, Eckart Meese, Andreas Keller

**Affiliations:** Helmholtz Institute for Pharmaceutical Research Saarland–Helmholtz Centre for Infection Research, 66123 Saarbrücken, Germany; Department for Clinical Bioinformatics, Saarland University, 66123 Saarbrücken, Germany; Institute of Human Genetics, Saarland University, 66421 Homburg, Germany; Department of Neurology, University Hospital of Saarland, 66421 Homburg, Germany; Institute of Virology and Center of Human and Molecular Biology, Saarland University, 66421 Homburg, Germany; PharmaScienceHub, Saarland University, Germany

**Keywords:** Parkinson’s disease, single-cell RNA sequencing, PBMCs, immune activation, PMA-Ionomycin, sex differences

## Abstract

Parkinson’s disease (PD) affects an estimated 6.4 million men and 5.3 million women worldwide, and still lacks a validated peripheral biomarker. Brain tissue is inaccessible in living patients, making peripheral blood an attractive alternative, but existing studies have profiled immune cells almost exclusively as static, resting-state snapshots that cannot reveal how those cells function under challenge. A longitudinal design, following patients over time, or applying a controlled stimulus to reveal functional differences invisible at rest, offers a more sensitive window onto disease-associated dysfunction, and sex, despite differential PD incidence and progression, is rarely treated as a primary variable. Here, using ex vivo PMA/ionomycin stimulation as a controlled functional challenge, we profiled 195k PBMCs from 84 samples of 14 PD patients and 14 controls by single-cell RNA sequencing across an activation time course (0h, 2h, 4h), stratified by sex throughout. Sex explained more transcriptional variance than disease status, and male and female PD patients showed largely divergent responses at rest that converged, by peak activation, on a discrete CD8+ effector memory T cell subpopulation (Tem-CD8). This subpopulation showed an exhaustion-consistent programme, coinciding with a failure to resolve AP-1 induction and a reduction in inferred intercellular communication. The PD peripheral immune phenotype is therefore better characterised as a activation-dependent response than a fixed resting-state signature, identifying Tem-CD8 exhaustion as a disease-associated, sex-convergent candidate for further study.

## Introduction

Parkinson’s disease (PD) is the fastest-growing neurological disorder worldwide. An estimated 8.5 million people were affected in 2019 (Luo et al. 2025), with projections suggesting this burden could double by 2040 (Dorsey, Sherer, et al. 2018; Dorsey, Elbaz, et al. 2018). The burden is consistently higher in men than in women (6.4 vs. 5.3 million prevalent cases worldwide in 2021) (Luo et al. 2025). Its defining motor features, including tremor, rigidity, and bradykinesia, emerge only after decades of subclinical neurodegeneration, by which point at least half of the dopaminergic neurons in the substantia nigra have already been lost (Kordower et al. 2013). Symptoms overlap with those of other movement disorders, and pathological changes precede clinical manifestation by years; as a result, diagnosis remains clinical, late, and imprecise.

Despite intense efforts, no blood-based peripheral biomarker has yet achieved sufficient sensitivity and specificity for routine diagnostic or stratification use. α-Synuclein seed amplification assays have recently transformed diagnostic performance in cerebrospinal fluid, the most validated specimen to date, while peripheral tissues including skin and blood show increasing but less standardised diagnostic utility (Bautista et al. 2026).

Among candidate sources of mechanistic insight and disease-associated signals, the peripheral immune system has attracted growing interest. It is accessible, responds dynamically to disease processes, and is increasingly implicated mechanistically in PD pathogenesis. Peripheral immune dysregulation is increasingly recognised as a component of PD, although its precise contribution to disease initiation and progression remains debated (Tansey et al. 2022; Tumpa et al. 2025). Genome-wide association studies reinforce this picture directly, identifying HLA and other immune-associated loci among PD genetic risk factors (Nalls et al. 2019). α-Synuclein-reactive CD4+ and CD8+ T cells are detectable in patient blood and post-mortem substantia nigra tissue, suggesting antigen-driven adaptive immune involvement (Sulzer et al. 2017; Brochard et al. 2009). Monocytes show elevated inflammatory gene expression in PD patients (Grozdanov et al. 2014). Pro-inflammatory cytokine profiles have also been reported across both bulk and single-cell studies. Single-cell resolution is particularly valuable here: it can resolve small, disease-relevant subpopulations that bulk profiling averages away, as recent PBMC-focused single-cell atlases and harmonized annotation resources illustrate (Grandke et al. 2025; Hoffmann et al. 2026). However, reported peripheral immune signatures vary considerably across cohorts, likely reflecting differences in disease stage, medication, technical platforms, and patient heterogeneity (Recinto et al. 2026). This variability has made it difficult to define a consistent, reproducible PD immune phenotype. Whether peripheral immune changes drive neurodegeneration or merely reflect it also remains unresolved.

A major limitation of existing studies is that they have largely characterised immune cells as static entities at rest, rather than as dynamic systems defined by their capacity to respond to stimulation (Xiong et al. 2024; Hong et al. 2025; Moquin-Beaudry et al. 2025). Chronic inflammatory exposure, persistent antigen stimulation, and age-associated immune remodelling have all been proposed to alter immune responsiveness in PD. This raises the possibility that dysfunction becomes most apparent during activation rather than at baseline. Consistent with this, α-synuclein-specific T cell reactivity itself follows a dynamic, disease-stage-dependent trajectory rather than a fixed resting state, rising before motor diagnosis and declining thereafter (Lindestam Arlehamn et al. 2020). Single-cell and genetic-association studies of resting PD PBMCs have identified altered or genetically-linked monocyte, NK-cell, and T-cell states (Xiong et al. 2024; Hong et al. 2025), but the activation capacity of these cells, their ability to mount and sustain a transcriptional response under challenge, has rarely been examined.

The use of ex vivo pharmacological stimulation to reveal disease- or state-associated immune phenotypes not apparent in resting cells is an established strategy. Functional perturbation assays have uncovered T cell deficits in chronic viral infection (Wherry et al. 2007), contraction of the naive T cell repertoire in immunosenescence (Franceschi et al. 1999), and sex-specific divergence in innate and adaptive immune responses following ex vivo stimulation (Fragiadakis et al. 2022). In PD specifically, Diener et al. demonstrated that CD2/CD3/CD28-mediated stimulation of isolated CD4+ T cells distinguishes PD patients from controls at the gene expression level in ways not detectable at rest (Diener et al. 2023). This provides direct precedent for the perturbation-based approach used here.

Profiling cells at multiple stages of a defined activation time course further enables discrimination between two kinds of response: early transient responses, which reflect acute signalling capacity, and late sustained programmes, which integrate transcriptional regulation, epigenetic state, and cellular fitness. This temporally contingent sequence of gene activation events is well established for lymphocyte activation broadly (Crabtree 1989). To our knowledge, no previous study has combined longitudinal activation profiling, single-cell transcriptomics, systematic sex stratification, and an inflammatory neurological comparator in PD. Such an approach may reveal sex-specific, activation-dependent immune phenotypes that are not apparent in resting PBMCs.

Ex vivo stimulation with PMA (Phorbol 12-myristate 13-acetate) and ionomycin bypasses antigen receptor engagement. It drives canonical lymphocyte activation programmes, including AP-1, NFAT, and downstream effector transcription, in a reproducible manner across donors. It does not measure immune competence in the broad sense, nor does it summarise antigen-specific responses. Rather, it provides a standardised perturbation that tests the maximal activation potential of the intracellular signalling programmes downstream of receptor engagement: phorbol ester and calcium ionophore act synergistically to bypass the receptor-proximal signalling steps that normally gate lymphocyte activation, engaging the PKC and calcium arms of the pathway directly instead (Truneh et al. 1985).

We included neurological control patients (predominantly ischaemic stroke, with a minority of other acute/subacute presentations; Methods) as an inflammatory neurological comparator. These patients provide a model of acute CNS injury accompanied by pronounced peripheral immune activation (Chamorro et al. 2012), but they lack the chronic neurodegenerative processes characteristic of PD. This lets us distinguish PD-specific immune alterations from generic CNS injury and neuroinflammatory responses.

Clinical immune-profiling data already distinguish male- and female-predominant peripheral inflammatory clusters in PD, with divergent clinical and central synucleinopathy correlates (Bovenzi et al. 2025). This further motivated a sex-stratified design. We hypothesised that ex vivo activation would uncover sex-stratified, activation-dependent immune phenotypes in PD that differ from healthy controls, and that single-cell time-course profiling would identify the cell types and programmes responsible. We collected PBMCs from 14 idiopathic Parkinson’s disease patients (PD), 5 age- and sex-matched healthy controls (HC), and 9 neurological control (NC) patients (clinical and demographic details in Supplementary Table 1), and stimulated them across a three-point time course (TP0: unstimulated; TP2: 2 h; TP4: 4 h). A separate pilot cohort of 4 young, healthy donors was used beforehand to validate the stimulation protocol and is not part of this main-cohort comparison.

## Results

### A time-resolved single-cell map of PBMC activation in Parkinson’s disease

To characterise the immune response to T cell activation in Parkinson’s disease, we collected PBMCs from 14 parkinson’s disease (PD) patients, 9 neurological control (NC) patients, and 5 healthy controls (HC) and stimulated them with PMA/Ionomycin across a three-point time course (TP0: unstimulated; TP2: 2 h; TP4: 4 h), yielding 84 samples in total (Figure 1 A–B). Neurological Control patients were included as an acute CNS inflammatory control to distinguish PD-specific immune phenotypes from generic neuroinflammatory responses.

**Figure 1:**
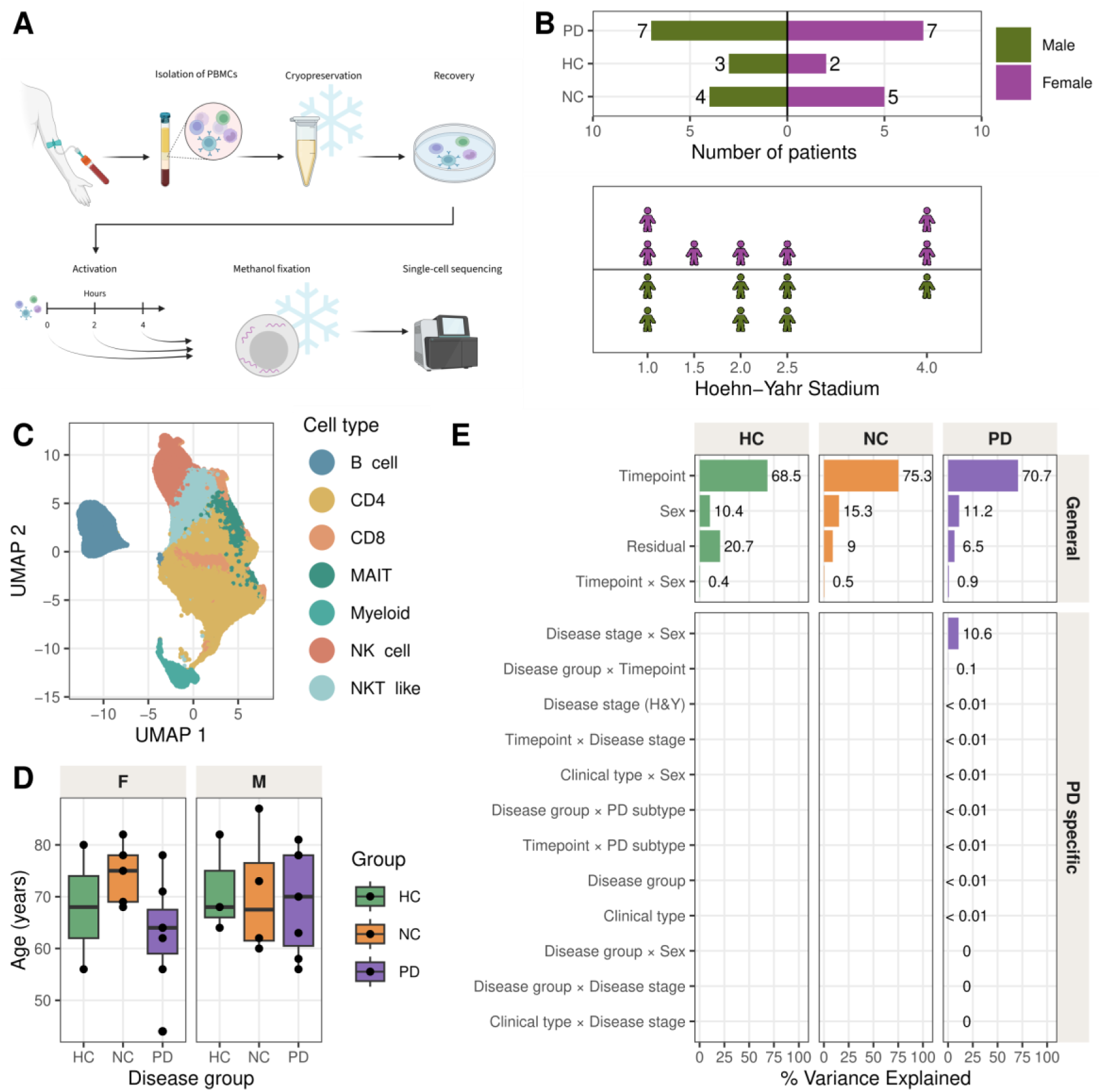
Study design and cohort overview. **(A)** Schematic overview of the PBMC PMA-Ionomycin activation protocol. Created in BioRender: Grandke, F. (2026) **(B)** Patient cohort composition by disease group (Parkinson’s Disease (PD), Neurological Control (NC), and healthy controls (HC)) and sex, and Hoehn-Yahr Stadium among PD patients. **(C)** UMAP representation of the dataset, coloured by broad cell type. **(D)** Age distribution stratified by disease group and sex. **(E)** PVCA results (pvcaBatchAssess threshold 0.6, combined-timepoint model) for general (sex, age, timepoint) and PD-specific factors (disease stage, disease group (CNS/PNS-first), clinical type (akinetisch rigide, aequivalenztyp, tremordominanztyp)).

After preprocessing and quality control (Supplementary Figure 1), we retained 195k cells across all samples (Figure 1 C). Cluster-based annotation yielded 23 fine-grained cell types across six lineages (CD4 T cells, CD8 T cells, NK cells, B cells, monocytes, and MAIT cells), comprising 147k T cells, 23k NK cells, 21k B cells, and 5k myeloid cells (Figure 1 C; Supplementary Figure 2). Cell type identity was confirmed by canonical marker gene expression and was consistent across stimulation timepoints (Supplementary Figure 2 A–B). Cell type proportions were stable across donors within each group, with expected inter-individual variation (Supplementary Figure 2 C). Disease groups did not differ significantly in age or sex (PD vs. HC: t-test p = 0.737 in females, p = 0.759 in males; NC vs. HC: p = 0.688 in females, p = 0.924 in males; Figure 1 D).

**Supplementary Figure 1:**
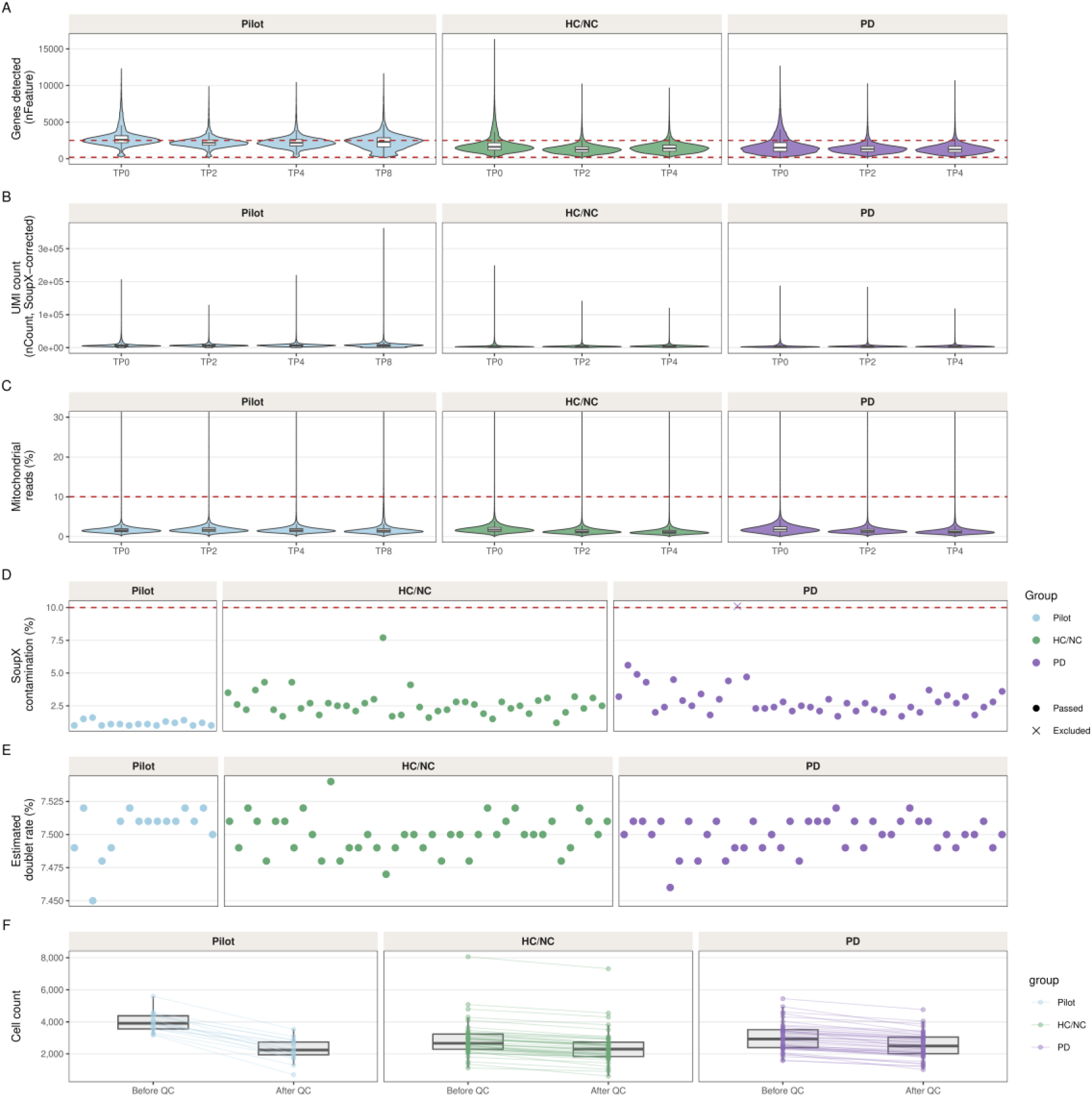
Single-cell RNA-seq quality control (unfiltered cells). **(A)** Genes detected (nFeature) per cell before filtering, grouped by cohort and timepoint. Dashed lines indicate the applied thresholds (200–2,500 genes). **(B)** SoupX-corrected UMI counts (nCount) per cell. **(C)** Mitochondrial read fraction (%) per cell; dashed line at 10% threshold. **(D)** SoupX ambient RNA contamination estimate per sample; × marks the excluded sample PDS_TP0 (10.1%; was re-sequenced and replaced). **(E)** Estimated doublet rate per sample (DoubletFinder). **(F)** Cell counts before and after QC filtering across samples.

**Supplementary Figure 2:**
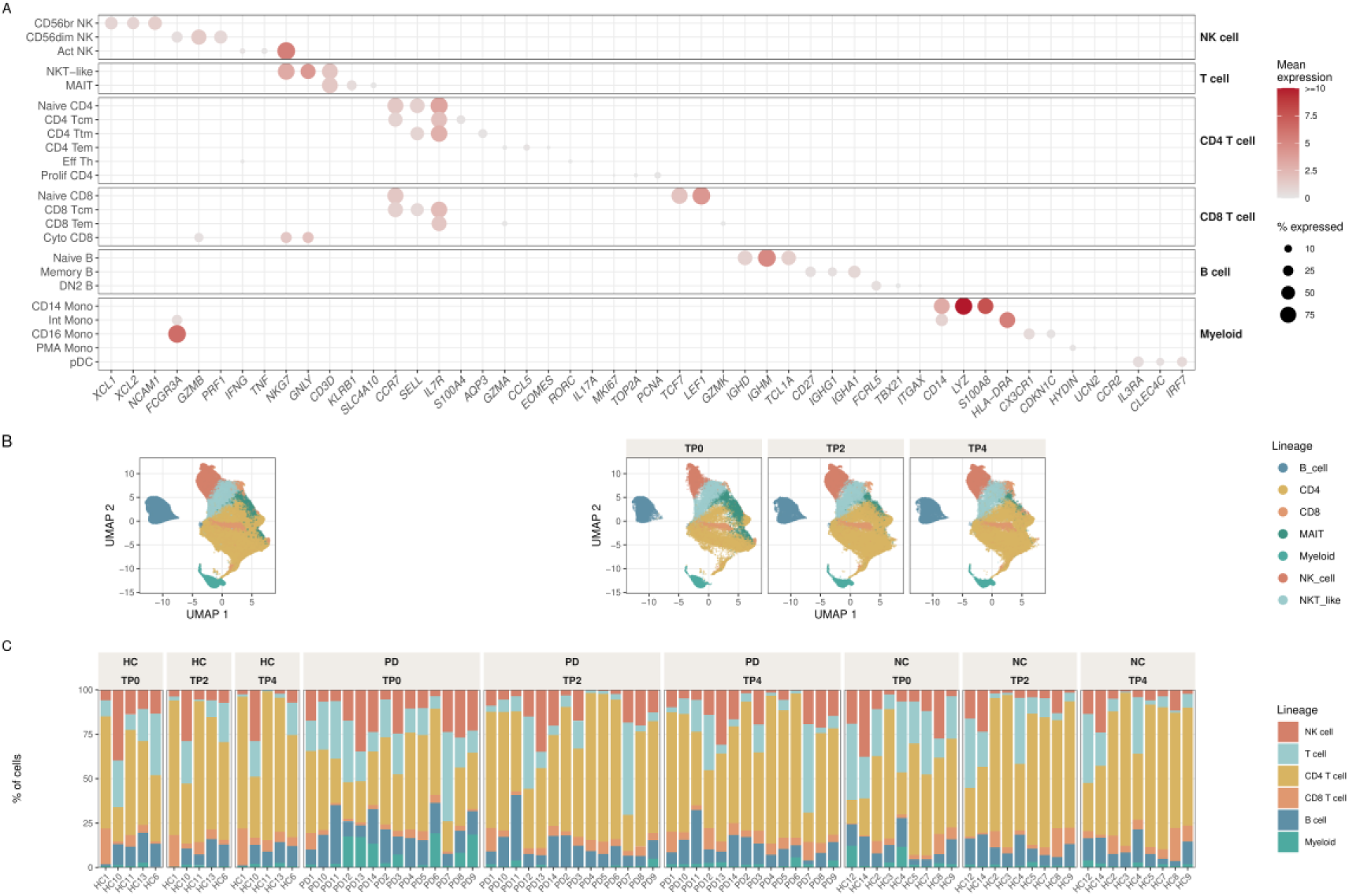
Cell type annotation validation. **(A)** Dot plot of canonical marker gene expression per cell type at TP0 (unstimulated). Dot size: percentage of cells expressing the gene; colour: mean log-normalised expression. **(B)** UMAP coloured by broad lineage annotation (left) and split by stimulation timepoint (right). **(C)** Stacked barplot of lineage-level cell type proportions per patient and timepoint, grouped by disease group.

Principal variance component analysis (PVCA) applied to the full pseudobulk expression matrix identified stimulation timepoint as the dominant source of transcriptional variance (70.7% in PD, 68.5% in HC), the expected signature of a strong, coherent stimulus (Figure 1 E). Sex accounted for 11.2% of variance in PD, while disease group (CNS-first vs. PNS-first) only contributed little to global transcriptional variance (< 0.01% in PD), indicating similar global immune activation capacity across PD subtypes.

### Technical variation (cryopreservation, cohort, storage duration) does not confound the activation response

Cryopreservation duration (days frozen) differed between HC and PD (median 43 vs. 230.5 days; Wilcoxon rank-sum test, two-sided, p = 0.001; single comparison, no multiple-testing correction applied), so we tested its downstream influence directly rather than assuming it away. We ruled out cryopreservation, freeze duration, and cohort as technical confounds (Figure 2). Viability was high in both cohorts (mean 86.5% pilot, 91.1% PD, 89% HC/NC), independent of cryopreservation duration (9–278 days; slope ≈ -0.0012, p = 0.931, n = 32), and freeze duration showed no association with the transcriptional profile genome-wide (BH-adj. p = 0.580/0.922/0.273 at TP0/TP2/TP4) or in a heat-shock/proteotoxic-stress panel (Lang et al. 2021) (p = 0.357/0.898/0.169, see “Methods: Module scores”). The groups’ unequal freeze durations were therefore not found to influence the transcriptional or viability outcomes reported here.

CD69 induction confirmed stimulation in pilot CD4+/CD8+ T cells (1.4% at rest to 96.2% at 2 h, 93.9% at 4 h, 93% at 8 h; Figure 2 D). Timepoint-pair fold-changes correlated closely (Pearson r > 0.4, adj. p <0.001; Figure 2 E), and the pilot UMAP reproduced the main cohort’s cell-type structure (Figure 2 F). TP8 was dropped: median cell count fell from 2604 at TP4 to 2115, consistent with activation-induced death beyond 4 hours. Mitochondrial read fraction stayed stable across the whole time course (median 1.6% at TP4 vs. 1.52% at TP8), so this is not a stress/apoptosis signature visible in that metric; genes detected per sample declined more directly (median 1958 at TP4 vs. 1739 at TP8), consistent with a genuine drop in per-cell library complexity by TP8 (Figure 2 G-H).

**Figure 2:**
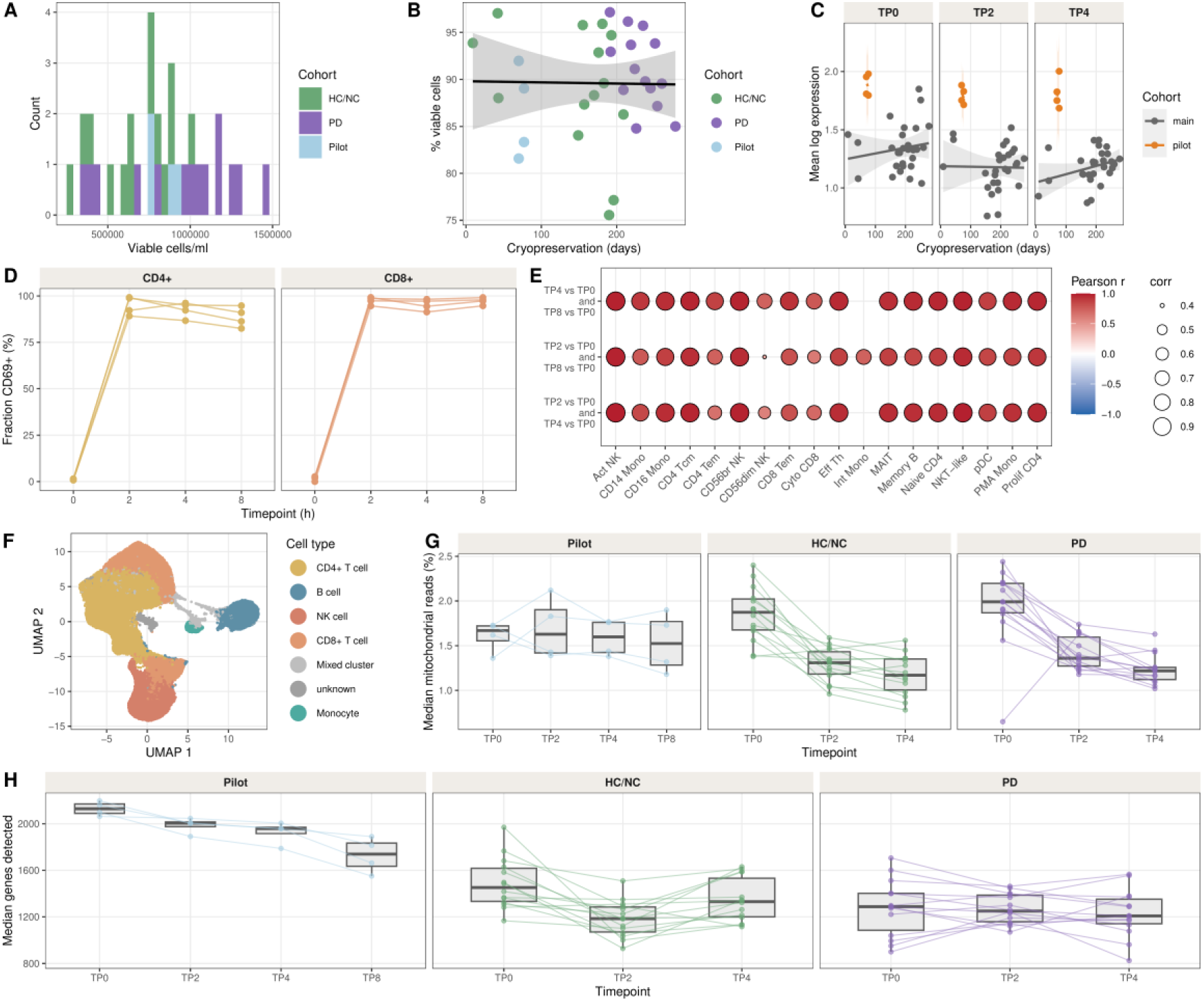
Pilot cohort validation of the PMA/Ionomycin time course (pilot: n=4 donors, 16 samples across TP0/TP2/TP4/TP8; main cohort: n=14 PD, n=9 NC, n=5 HC). **(A)** Distribution of viable cell counts per ml across pilot and main cohort samples. **(B)** Viable cell proportion vs. cryopreservation duration (linear regression; shaded band: 95% confidence interval of the linear fit). **(C)** Per-sample overall expression level (mean log-transformed pseudobulk expression) vs. cryopreservation duration, by timepoint and cohort (shaded band: 95% confidence interval of each per-cohort linear fit). **(D)** FACS: fraction of CDcS+ cells within CD4+ and CD8+ T cells at TP0, TP2, TP4, and TP8. **(E)** Fold-change correlation heatmap between pilot timepoint pairs; significant (two-sided Pearson correlation, adj. p < 0.05, BH adjustment) correlations have black borders. **(F)** UMAP of pilot cohort scRNA-seq data coloured by manual annotation. **(G)** Median mitochondrial read fraction (%) per sample across timepoints. **(H)** Median genes detected per sample across timepoints.

Sex concordance of per-cell-type log2 fold-changes (Pearson r; paired, patient-blocked limma-voom design) across HC donor pairs was high at TP0-TP2 (median r = 0.92) and declined moderately by TP4 (median r = 0.56), plausibly reflecting sex differences in innate immune responsiveness (Supplementary Figure 3 A). Concordance between the pilot cohort (n=4 donors, all female, all 22 years old) and older HC females was more moderate (median r = 0.61 at TP0-TP2, 0.34 at TP4), likely reflecting inter-individual variability rather than age, since this pairs a young, homogeneous group against an older, more heterogeneous one.

Cross-disease-group concordance of per-cell-type log2 fold-changes (paired, patient-blocked design) between HC, PD, and NC was broadly conserved across the early (TP0-TP2) and late (TP2-TP4) phases (Supplementary Figure 3 A): mean Pearson r at TP0-TP2 was 0.895 (HC vs PD), 0.894 (HC vs NC), and 0.873 (PD vs NC), declining to 0.584, 0.641, and 0.639 at TP2-TP4. This late-phase decline is led by EffectorCytotoxic CD8 (mean cross-group r = 0.05 in the TP2-TP4 window, the lowest of any cell type); PMA-induced monocytes likely also contribute, a population virtually absent at baseline (0.8% of myeloid cells at TP0 vs. 55.6% at TP4), so its fold-changes reflect baseline abundance rather than divergent transcriptional programmes. Larger activation signatures were also more concordant across groups (Supplementary Figure 3 B; slope = 0.05 per log₂FC unit, R² = 0.36).

Together, these results show a comparable, cell-type-resolved activation response across disease group and sex. The age comparison, limited by the single-donor pilot design, shows moderate TP4 concordance but no systematic age-specific programme. Quantitative differences between groups, addressed below, therefore reflect biological rather than technical divergence.

**Supplementary Figure 3:**
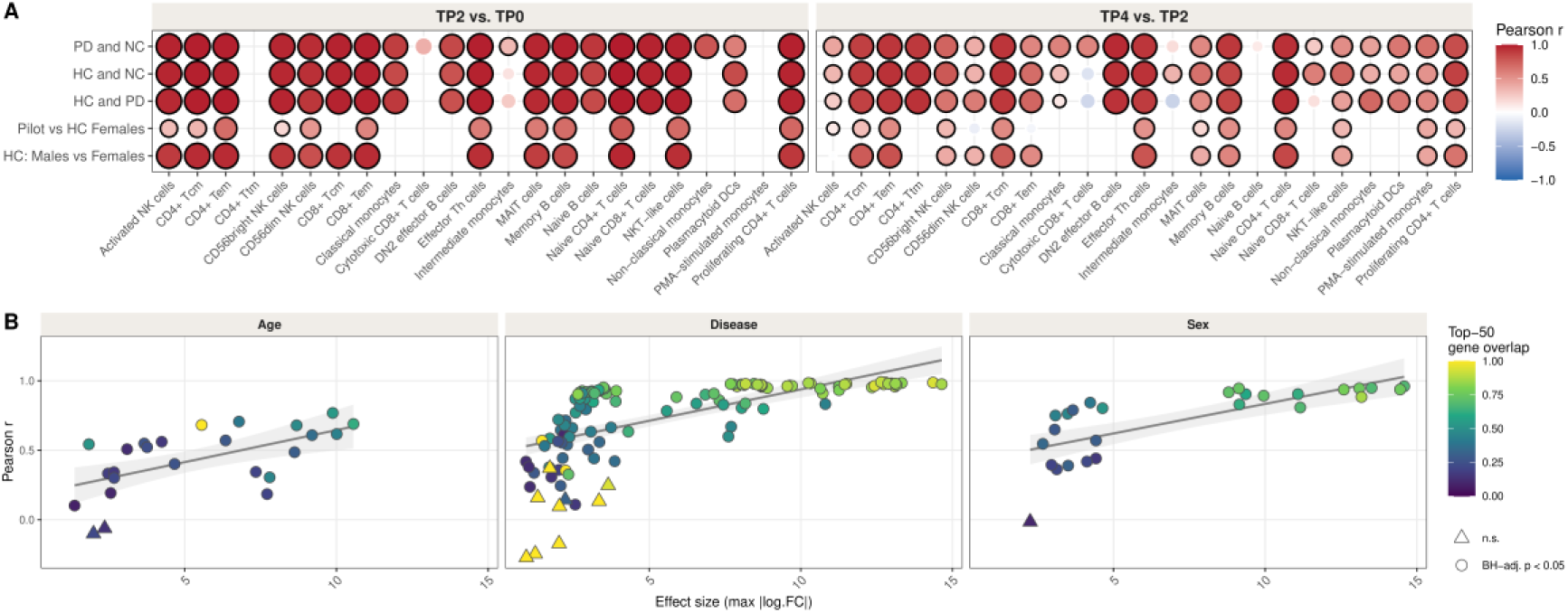
Cross-subgroup concordance of the PMA/Ionomycin transcriptional activation response. (A) Pearson r (two-sided test, BH-adjustment) of gene-level log2 fold-changes (paired, patient-blocked limma-voom design) per cell type between five subgroup comparisons (rows: HC male vs. female donor pairs, n=3 vs. n=2; pilot cohort vs. HC females, n=4 vs. n=2; HC vs. PD, n=5 vs. n=14; HC vs. NC, n=5 vs. n=9; PD vs. NC, n=14 vs. n=9) and two activation phases (columns). black border = significant. (B) Pearson r (two-sided test, BH-adjustment) between subgroup log2FC vectors vs. maximum absolute log2FC per cell type and comparison, faceted by Sex/Age/Disease subgroup; shaded band: 95% confidence interval of the linear fit (grey line) across all points.

### HC Activation Context: The Healthy Immune Response

PMA/Ionomycin stimulation induces a two-phase transcriptional response in healthy PBMCs. The early phase (TP0→TP2) is dominated by immediate-early genes broadly upregulated across lymphoid and myeloid compartments, including CSF2, IL2, IFNG, CCL4L2, XCL1 (Supplementary Figure 4 A); the late phase (TP2→TP4) is narrower and more cell-type-specific. Hallmark pathway enrichment follows the same two-phase pattern (Supplementary Figure 4 B). This two-wave structure is consistent across lymphoid lineages and is the reference frame against which PD-specific deviations are interpreted throughout.

PMA/Ionomycin also activates three signalling axes: Ca²⁺/NFAT, PKC/NF-κB, and PKC/AP-1 (Supplementary Figure 4 C, see “Methods: Module scores”). PKC/AP-1 showed the broadest activation at TP0→TP2; Ca²⁺/NFAT and PKC/NF-κB were most active in T and MAIT cells, matching their established role in TCR signalling. Monocyte arm-gene coverage was comparatively sparse and high-variance, with PKC/NF-κB trending up and PKC/AP-1 trending down.

PD-vs-HC differences in arm activation at TP0→TP2 were modest (Supplementary Figure 4 D): most cell type × arm combinations differed by less than 0.5 log₂FC from HC, and none of the larger differences survived multiple-testing correction, so these comparisons remain exploratory.

**Supplementary Figure 4:**
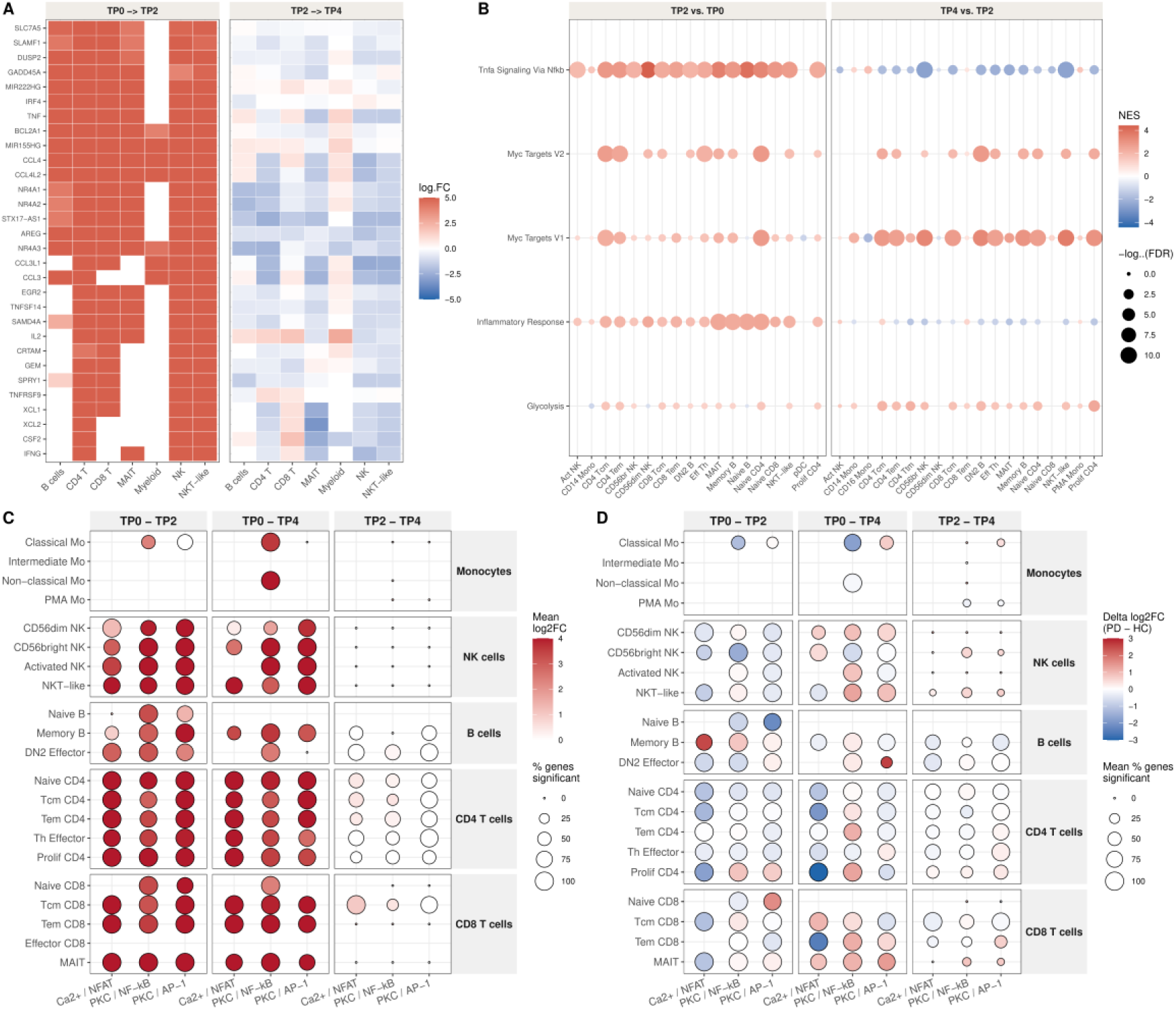
The activation programme in healthy PBMCs: transcriptional waves and signalling arms (HC, n=5; panel D additionally compares against PD, n=14). **(A)** Heatmap of the 30 most broadly differentially expressed genes (paired, patient-blocked limma-voom; BH-adjusted adj. p < 0.05 & |log₂FC| > 0.5 for the underlying significance calls) at early (TP0→TP2) and late (TP2→TP4) activation in HC (genes selected by breadth × effect size with a minimum expression filter). **(B)** Top 3 Hallmark pathways per phase by normalised enrichment score (NES; preranked GSEA, gseapy). **(C)** PKC/AP-1, PKC/NF-κB, and Ca²⁺/NFAT signalling-arm scores in HC across three timepoint comparisons (paired, patient-blocked limma-voom; BH-adjusted adj. p < 0.05). **(D)** PD vs. HC difference in signalling arm activation (PD mean log₂FC minus HC mean log₂FC).

Cell type proportions shifted substantially with activation (Figure 3 A-C). A monocyte population virtually absent at rest emerged progressively by TP4, hereafter “PMA-induced monocytes,” increasing strongly in all three disease groups (DA Estimate: 2.42 and 3.07 in HC, 1.3 and 2.8 in PD, 2.05 and 3.17 in NC; all adj. p < 0.05), an internal validation of the stimulation protocol shared by CD8+ Tcm/Ttm and DN2 effector B cell increases and Naive B, Naive CD8+, MAIT, and CD56dim NK declines (Figure 3 C). A smaller set was PD-specific: Intermediate monocytes (Estimate: -1 and - 1.08, adj. p = 0.045 and 0.031), Non-classical monocytes (Estimate: -1.47 and -1.18, adj. p = 0.009 and 0.031), and NKT-like cells (Estimate: -0.71 and -0.6, adj. p = 0.004 and 0.012) declined significantly in PD but not in HC or NC.

**Figure 3:**
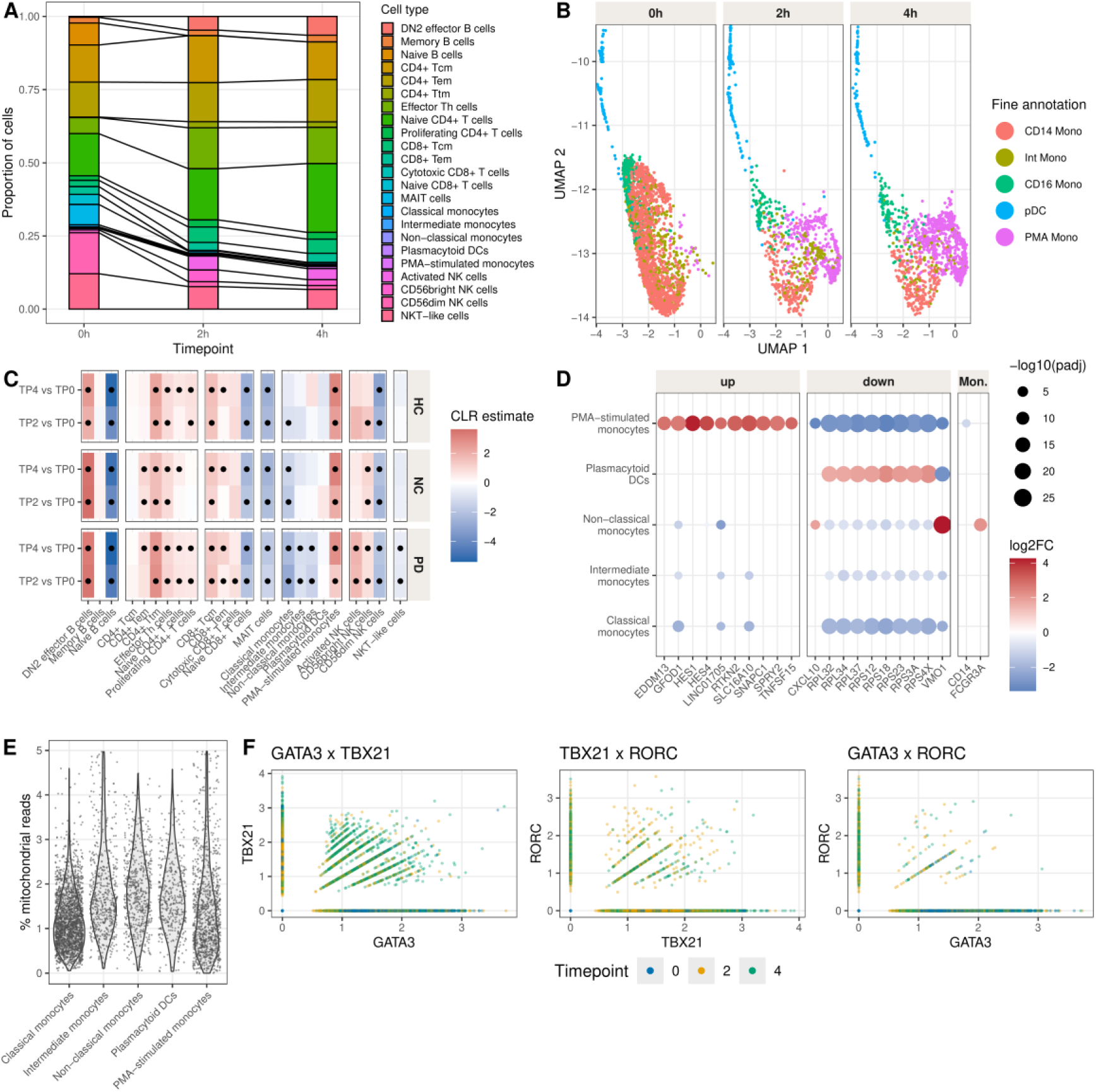
Activation-induced compositional shifts across PBMC lineages (n=14 PD, n=S Neurological Control, n=5 HC; panel A is HC-only). **(A)** Cell type proportions at TP0, TP2, and TP4 in HC. **(B)** UMAP of myeloid cells across all groups (HC, PD, NC), faceted by timepoint. **(C)** Longitudinal differential abundance (CLR-transformed proportions, patient-blocked linear model, BH-adjusted); filled dots = significant (adj. p < 0.05). **(D)** PMA-induced monocyte marker genes (unpaired pseudobulk limma-voom, one-vs-rest per myeloid subtype, BH-adjusted), grouped into three facets: Top 10 most significantly up- and top 10 most significantly downregulated genes and CD14/FCGR3A as canonical monocyte identity markers. **(E)** Percentage of mitochondrial reads per cell across myeloid subtypes, including PMA-induced monocytes. **(F)** Pairwise co-expression of the lineage-associated transcription factors GATA3, TBX21, and RORC in T cells.

Differential expression against other myeloid subtypes characterised the PMA-induced monocyte state, most strongly HES1 (log2FC = 4.17, adj. p = 3.5e-18) and HES4 (log2FC = 3.57, adj. p = 5.8e-15) upregulated, and, among the downregulated genes, VMO1 (log2FC = -3.28) and CXCL10 (log2FC = -3.22) alongside a broad reduction in ribosomal protein genes consistent with lower translational capacity (80 genes; Figure 3 D); because this test pools cells across all three timepoints and the PMA-induced monocyte cluster is only prevalent at TP2/TP4, these markers cannot be fully separated from general late-timepoint activation genes. Mitochondrial RNA content remained normal (Figure 3 E), consistent with a reversible activation state rather than de novo differentiation. A subset of T cells transiently co-expressed TBX21 and RORC during early activation without stable terminal polarisation (Figure 3 F): the double-positive fraction rose from 0.015% at TP0 to 0.369% at TP2 (OR = 25.1 vs. TP0) and remained at a similar odds ratio of 25.1 by TP4 rather than increasing further, whereas the TBX21-only fraction increased steadily and more than the double-positive fraction (3.31% at TP0 to 31.35% at TP4), a pattern consistent with transient co-expression rather than progressive double-positive polarisation.

## PD leaves an activation-independent transcriptional imprint at rest

Parkinson’s disease alters peripheral immune transcription even without exogenous stimulation (Grozdanov et al. 2014). We characterise this resting-state signal as gene–cell type pairs consistently dysregulated across both sexes and all three timepoints.

Due to sample size limitations, we prioritised cross-timepoint reproducibility over single-timepoint significance, retaining a gene–cell-type pair only if it showed a consistent effect size and direction at TP0, TP2, and TP4. Naive single-timepoint pseudobulk comparisons (PD vs. HC) indeed yield few DEGs at any timepoint alone (2/28/6 at TP0/TP2/TP4), consistent with this expectation.

We asked whether any genes show PD-associated deregulation independent of activation state, using an effect-size consistency criterion (|log2FC| ≥ 0.5, same direction at TP0, TP2, and TP4; limma-voom pseudobulk), run separately by sex (male PD vs. male HC; female PD vs. female HC) to remove sex-composition confounds. Gene–cell-type pairs were classified as concordant (both sexes, same direction, the primary signature), discordant (both sexes, opposing directions), or sex-specific (one sex only) (Figure 4 A).

**Figure 4:**
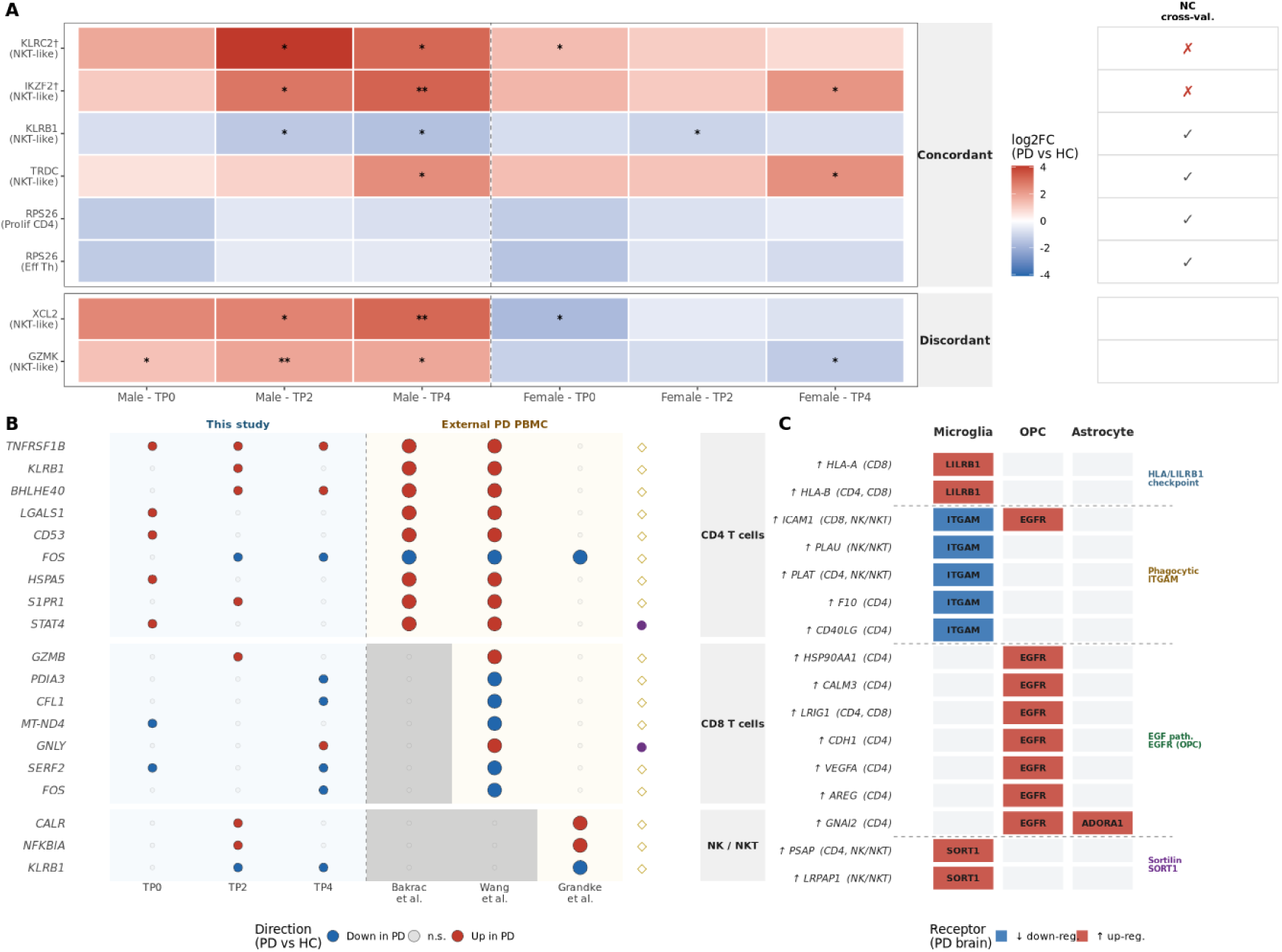
The PD peripheral transcriptional signature at rest: sex-stratified, cross-study validated, and linked to brain gene expression. (A) Sex-stratified, activation-independent PD transcriptional signature (|log2FC| ≥ 0.5, same direction at TP0, TP2, and TP4) in male and/or female PD vs HC comparisons (PD: 7M/7F; HC: 3M/2F); * p < 0.05 nominal, ** p < 0.05 BH-adjusted (limma-voom pseudobulk moderated t-test, two-sided). Right strip: NC vs HC cross-validation (mixed sex, Concordant category only); † on a gene label marks a pair not replicated in NC vs. HC. (B) Cross-study consistency (This dataset + Bakrač/Wang for CD4, Wang for CD8, Grandke et al. for NK (Grandke et al. 2025))of PD-associated CD4/CD8/NK T cell genes, computationally selected genome-wide (two-step criterion - own-data significance plus agreement, then significance in every well-powered external comparator for that lineage). Full set in Supplementary Figure 5; shown here is the literature-supported subset, see Supplementary Table 2. ● = prior PD/neurodegeneration literature; ◇ = adjacent literature. (C) Exploratory ligand-receptor bridging between PD-upregulated T/NK ligands (PBMC cohort) and brain DEGs from three independent PD snRNA-seq studies (Smajić et al. SN midbrain; Martirosyan et al. SNpc; Zhu et al. prefrontal cortex; pairs shown only if confirmed in ≥2). Ligand-receptor pairs from OmniPath (ligrecextra).

We identified 6 concordant gene–cell-type pairs across 5 unique genes (Figure 4 A), plus 2 discordant pairs. The concordant set is dominated by NKT-like cells and reveals two themes. The first is a dysregulated NK/NKT programme: IKZF2 (HELIOS) is consistently elevated (log2FC ≈ 1.11/1.54 in males/females at TP0; the only gene reaching BH-significance, p_adj = 0.048–0.009), KLRC2 (NKG2C) and TRDC (T cell receptor delta constant region, co-expressed with NK receptors in NKT cells) are both up, and KLRB1 (CD161) is down, a shift in activating/inhibitory receptor balance. The second is a translational axis: RPS26 is consistently downregulated across 2 CD4+ cell types in both sexes. Only IKZF2 reaches BH-significance; KLRB1, KLRC2, and TRDC reach nominal significance, and RPS26 is detectable only through the consistency criterion, these results should thus be read cautiously.

Cross-validating against Neurological Control patients (same consistency criterion) separates PD-specific biology from generic neuroinflammation: 4 of 6 concordant pairs (KLRB1, RPS26, TRDC) were also FC-consistent in Neurological Control vs HC, indicating a shared neuroinflammatory response rather than PD-specific biology (Figure 4 A, right strip). IKZF2, by contrast, distinguishes PD from both healthy and inflammatory-neurological-disease controls: it is elevated in PD relative to NC (log2FC ≥ 1.4) as well as relative to HC. KLRC2 shows a similar absence in Neurological Control at TP0 but not the same sustained PD > Neurological Control enrichment: its PD-vs-NC log2FC falls from 1.66 at TP0 to -0.2 by TP4.

To place these activation-independent findings in broader context, we asked two follow-on questions: which PD-associated signals replicate in independent blood studies, and whether the T and NK cell ligands elevated in PD could hypothetically engage brain immune cell states. We compared DEGs across CD4 T cells, CD8 T cells, and NK/NKT cells against three published PD single-cell datasets (Bakrac et al. (Meglaj Bakrač et al. 2026) [CD4-only], Wang et al. (Wang et al. 2021) [T cells], Grandke et al. (Grandke et al. 2025)), then performed an exploratory ligand-receptor analysis against brain snRNA-seq DEGs from three PD brain studies.

### Cross-study consistency of PD-associated T cell gene expression

Cross-study comparison identifies PD-associated transcriptional changes that replicate across studies and lineages (Figure 2 B, Supplementary Figure 5). A gene enters this analysis if it is nominally significant in our own data and reaches significance, in the same direction, in every well-powered external cohort available for its lineage.

Among CD4 T cell genes passing this computed selection, FOS is the cleanest confirmation: consistently down-regulated across Bakrac et al. and Wang et al., and independently confirmed as a significant down-DEG in an external PD scRNA-seq dataset. TNFRSF1B, LGALS1, HSPA5, and S1PR1 also replicate on this cross-study basis, each with genuine but indirect Parkinson’s-relevant literature: TNFRSF1B has weak, inconsistent protein-level evidence; Galectin-1 (LGALS1) and GRP78/HSPA5 have documented roles in rodent PD models rather than human PD immune cells, with a direct test of GRP78 in PD patient CD4+ T cells finding no significant difference from controls; and S1PR1’s link is limited to a pharmacological MPTP-mouse-model study. All four are therefore treated as neurodegeneration-adjacent rather than direct PD literature support (Supplementary Table 2). Two further genes, TNFAIP3 and CD5, replicate at least as strongly as any of these but no supporting PD-immune-cell literature was found for either.

In CD8 T cells, GNLY (up) carries the strongest support: enhanced expression in CD8 TEM/TEMRA has been directly reported by an independent PD scRNA-seq study. Five further replicating genes (GZMB, PDIA3, CFL1, MT-ND4, SERF2) have mechanistic links to α-synuclein aggregation or PD-associated cytotoxicity, though each link is indirect: from brain tissue, in vitro/invertebrate assays, or, for MT-ND4, the opposite direction in the one PBMC-level dataset available. In NK/NKT cells, KLRB1, NFKBIA, and CALR replicate in Grandke et al., the only dataset available for this lineage; none of the three has direct PD-immune-cell literature support beyond that replication. NDUFA11 replicates computationally in CD8 T cells but, like TNFAIP3 and CD5 above, has no supporting literature at all: mitochondrial complex I dysfunction is a well-established PD mechanism in dopaminergic neurons, but no NDUFA11-specific or T-cell/PBMC evidence was found, so it too is shown only in the full computed list.

**Supplementary Figure 5:**
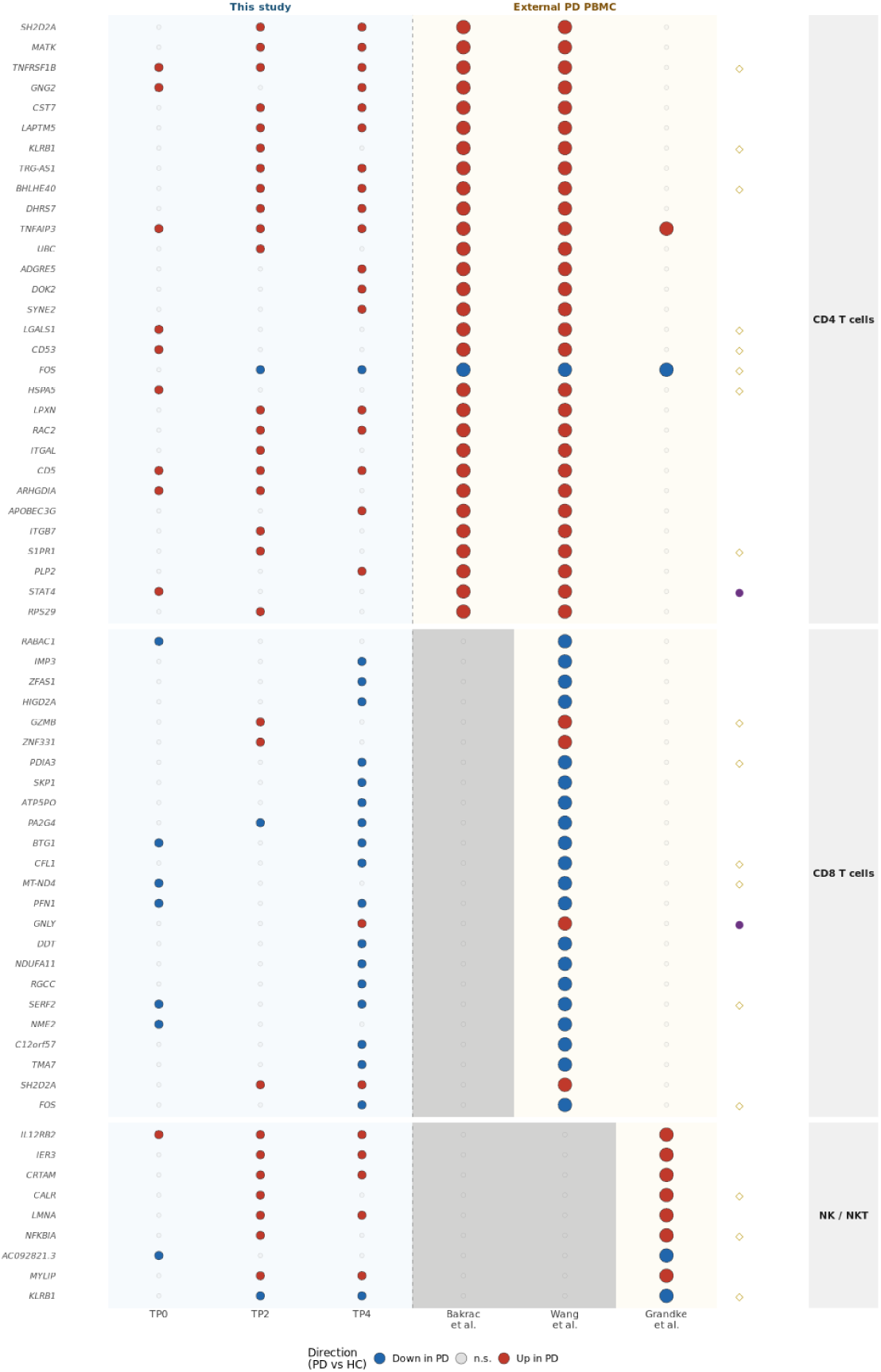
Full computed cross-study candidate pool (63 genes: 30 CD4, 24 CD8, 9 NK/NKT), before the literature filter applied for the main-text subset (Figure 4 B; Methods). Every gene here independently passes: (1) nominally significant in our own data at one timepoint, with at least one other timepoint agreeing in direction; (2) that direction significant in every well-powered external comparator available for its lineage. Columns and symbols as in Figure 4 B. This is the full, unfiltered result of the computed selection rule: nothing shown in the main-text figure was chosen from outside this list, and nothing here was excluded from the main figure for any reason other than the stated literature criterion (full status per gene in Supplementary Table 2 for the main-figure subset; genes shown here only were not literature-annotated). Full data in Supplementary Data.

### Exploratory cross-tissue ligand-receptor analysis

To test whether PD-associated changes in peripheral T cells might hypothetically influence brain immune cell states, we performed an exploratory ligand-receptor bridging analysis: PD-upregulated T cell genes (p < 0.05) were filtered to 109 known extracellular ligands (OmniPath ligrecextra) and their cognate receptors checked against brain snRNA-seq DEGs from three independent PD or Lewy body disease studies (Smajić et al. (Smajić et al. 2022); Martirosyan et al. (Martirosyan et al. 2024); Zhu et al. (Zhu et al. 2024)).

Of 109 candidate ligands, 66 had a cognate receptor among brain DEGs; 16 of these, forming 5 distinct receptor-by-brain-cell-type pairings, recurred across ≥2 studies (Figure 4 C). Because the datasets are non-matched and cross-sectional, none of the following can show that a peripheral ligand and a brain receptor ever actually encounter each other in the same tissue, let alone the same individual; every pairing below is an untested hypothesis, not evidence of an interaction. HLA-A/B pairs, in the OmniPath database, with the inhibitory receptor LILRB1 (up in PD microglia), consistent with elevated peripheral HLA-A/B engaging an inhibitory checkpoint on PD microglia. Coagulation and adhesion ligands (ICAM1, PLAT, PLAU, F10, CD40LG) pair, per OmniPath, with the phagocytic integrin ITGAM (down in PD microglia), consistent with a role in modulating microglial phagocytic capacity. EGF-pathway ligands (LRIG1, VEGFA, AREG, CDH1, CALM3, HSP90AA1) pair with EGFR (up in PD OPCs), pointing to a possible peripheral-to-OPC EGF signalling axis. GNAI2 additionally pairs with ADORA1 (up in PD astrocytes), suggesting adenosinergic signalling to astrocytes. PSAP pairs with microglial SORT1, echoing the sortilin-progranulin axis in GRN-linked neurodegeneration, though that axis concerns progranulin specifically.

### Disease severity scales with a Tcm-CD4 NR4A programme

Because all PD patients were treated, and treatment burden may track with disease progression, we first assessed this confound before correlating gene expression with Hoehn-Yahr Stadium.

14 of 14 patients were on at least one dopaminergic medication (Supplementary Figure 6), and medication load correlated positively with Stadium (Spearman rho = 0.8, p < 0.001), an unavoidable collinearity in this fully-treated cohort.

**Supplementary Figure 6:**
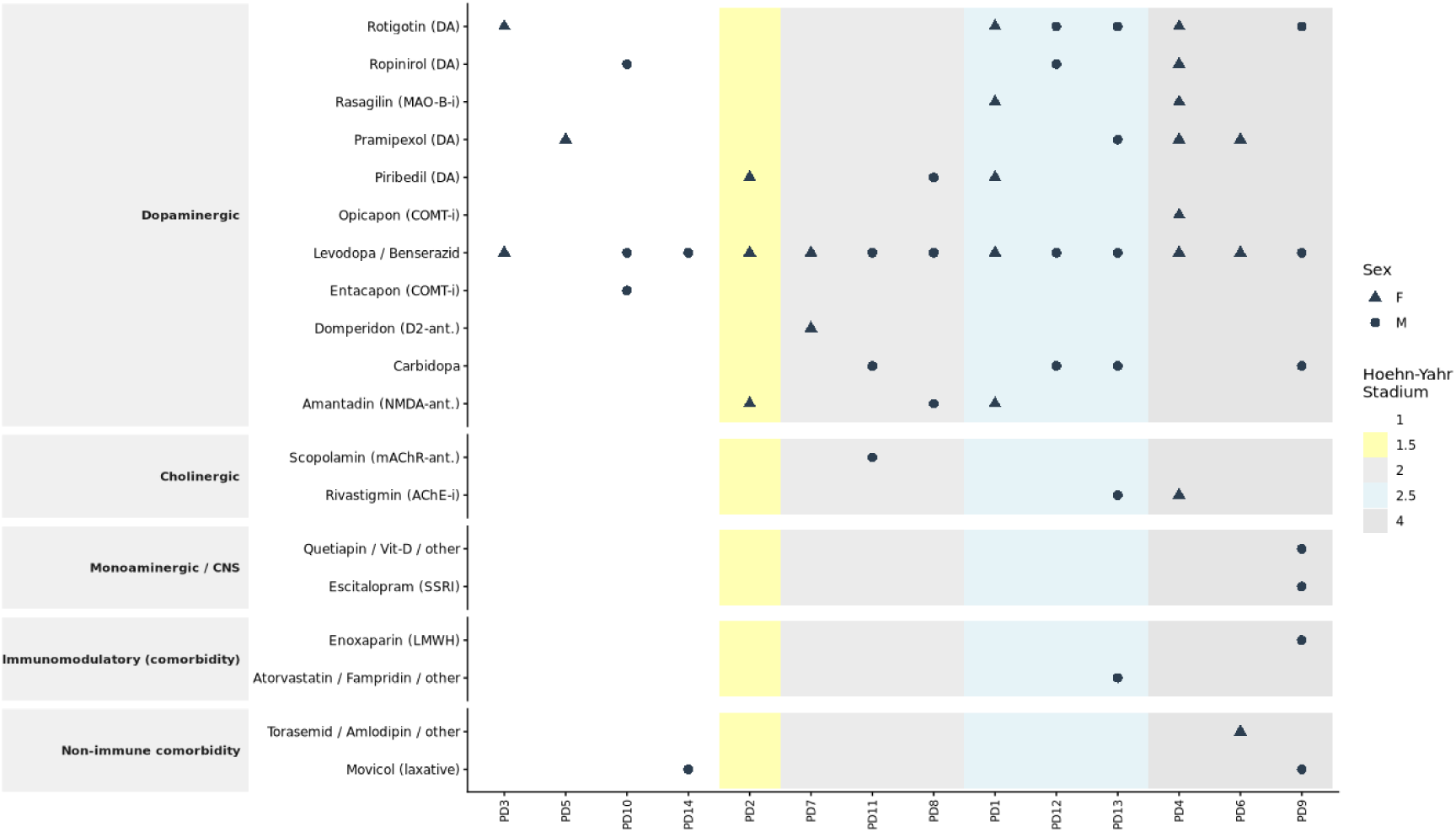
Medication profile of PD patients (n = 14). Medication grouped by pharmacological mechanism.

Independent of sex, we asked whether disease severity correlates with transcription across the three activation timepoints. The PD cohort’s sex and Hoehn-Yahr distributions were balanced (Wilcoxon p = 1; Figure 1 B). Per-gene Pearson correlations within PD patients (n ≥ 10; |r| > 0.5, p < 0.05 at each timepoint) showed most severity associations are activation-state-dependent: only 20 gene-cell type pairs held a consistent direction across TP0, TP2, and TP4 (Figure 5 A), dominated by Tcm-CD4 (14 of 20, all positive). Its flagship member, NR4A2, correlates with Stadium at every timepoint (r = 0.55/0.74/0.74; Figure 5 B), the same nuclear receptor family as NR4A1/NR4A3, later the top cross-sex-concordant genes in Tem-CD8. This apparent stability across timepoints masks a sex asymmetry: the TP0 leg is near-null in males (r = 0.221) and driven almost entirely by females (r = 0.774), whereas TP2 and TP4 are concordant in both sexes (male r = 0.812/0.544, female r = 0.697/0.858). BBC3 and GLUL complete the set; all four genes’ correlations persist after controlling for medication load (6 of 8 tested gene-timepoint correlations retain most of their effect size once medication load is controlled for; the remainder attenuate substantially, consistent with a partial medication-dose contribution specifically for those findings).

**Figure 5:**
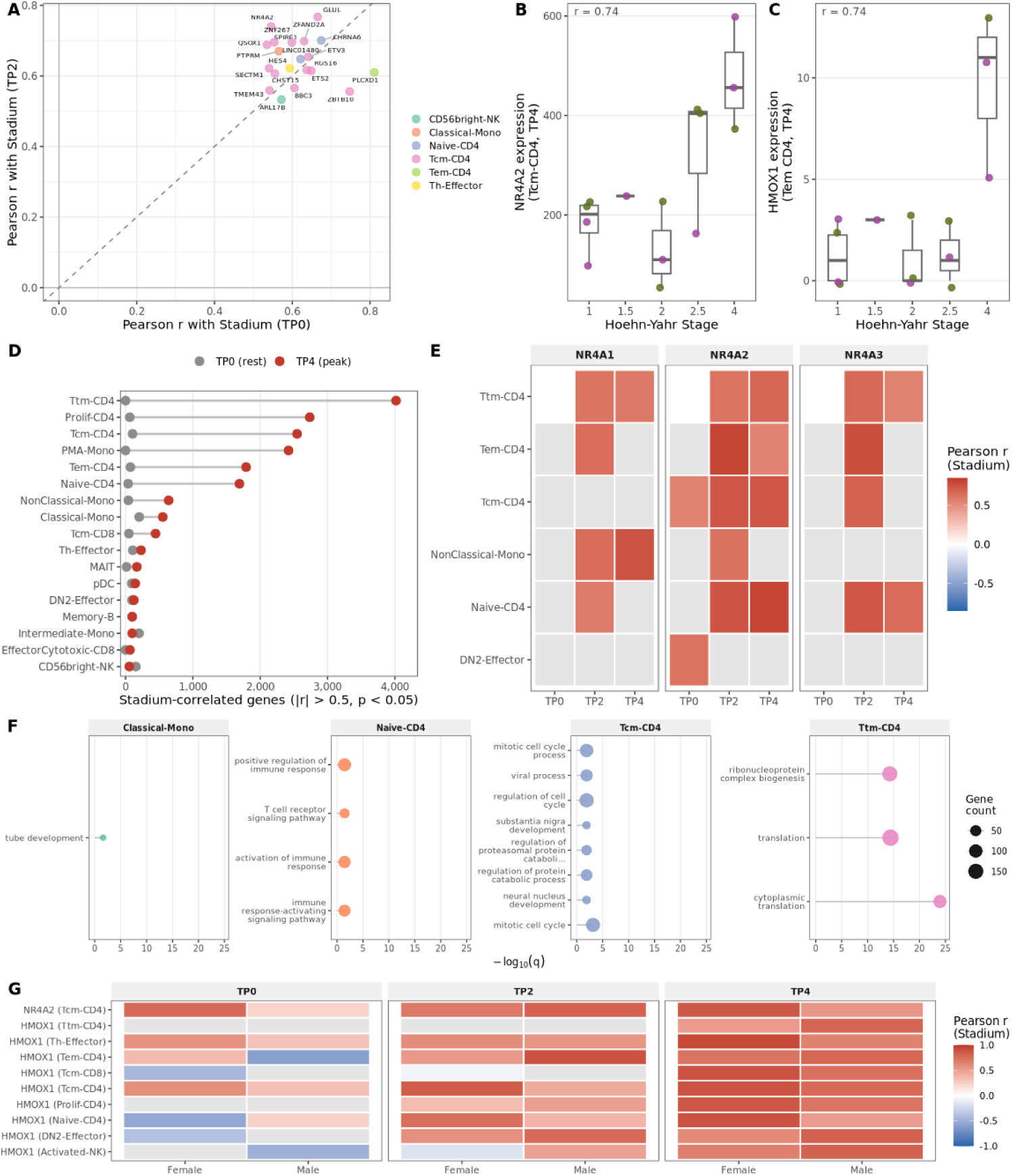
Hoehn-Yahr Stadium correlations with the PBMC transcriptome across activation timepoints (Stadium distribution across male and female PD patients shown separately in Figure 1 B). (A) The 20 gene–cell type pairs with consistent directional Stadium correlation across all three timepoints (TP0, TP2, TP4; Pearson correlation, two-sided; |r| > 0.5, p < 0.05 nominal at each timepoint independently; n ≥ 10 per timepoint). (B) NR4A2 expression per patient at TP4 (Tcm-CD4) by Stadium. (C) HMOX1 expression per patient at TP4 by Stadium, in Tem CD4. (D) Per-cell-type count of Stadium-correlated genes (Pearson correlation) at rest (TP0) and peak activation (TP4); cell types with ≥ 50 correlated genes at TP4 shown. (E) NR4A family (NR4A1/2/3) Stadium correlations; tiles show Pearson r only where significant (two-sided; |r| > 0.5, p < 0.05 nominal). (F) GO Biological Process over-representation analysis of genes positively correlated with Stadium at both TP2 and TP4 (Pearson correlation, two-sided; r > 0.5, p < 0.05 nominal; background = 8043 genes); top 3 terms per cell type (q < 0.05, BH-adjusted). (G) Sex-stratified Stadium correlations for the genes discussed individually in the text (Pearson correlation, two-sided, r within male/female PD patients, n = 6-7 per sex) per timepoint.

NR4A2/NURR1’s positive correlation with Stadium at rest (TP0: r = 0.55) is not directly comparable to prior reports of decreased NURR1/NR4A2 mRNA in resting peripheral lymphocytes from PD patients versus healthy controls (Le et al. 2008; Liu et al. 2012): those studies used a case-control design in mixed lymphocyte populations, whereas this is a within-PD, severity-scaled correlation specific to Tcm-CD4. A gene can be lower in PD than in HC overall while still scaling positively with severity among PD patients themselves.

A second individual finding at TP4: HMOX1, a PD risk gene and the principal cytoprotective oxidative-stress response, is the most broadly Stadium-correlated gene overall (9 cell types, strongest in Tem CD4 at r = 0.74; Figure 5 C), consistent with an activation-revealed oxidative-stress gradient tracking motor severity. Beyond these individual genes, the Stadium-transcriptome relationship is nearly invisible at rest and emerges almost entirely under activation (Figure 5 D): 1701 gene-cell type pairs correlate with Stadium at TP0, versus 17911 at TP4 (a 11-fold expansion, nominal significance only), most extreme in CD4+ T cells; NR4A1/NR4A3 show the same pattern, absent at rest and detectable only under activation (Figure 5 E). GO enrichment on the TP2/TP4-correlated genes found significant terms in 4 of 10 cell types tested (Figure 5 F), with Tcm-CD4’s top term pointing to a proliferative programme alongside the exhaustion and apoptosis markers above. Sex-stratified correlations for these genes (Figure 5 G) are consistent with the NR4A2 pattern described above.

Because this comparison uses nominal, uncorrected significance, we repeated it under genome-wide and per-cell-type BH correction: no pair survives genome-wide correction at any timepoint, and per-cell-type correction leaves only 4 surviving pairs, all at TP2 and none of them NR4A2, BBC3, or GLUL. This confirms the three-timepoint-consistent signature’s evidence rests on an independent criterion, a consistent direction at nominal p < 0.05 across all three timepoints, rather than on single-timepoint significance surviving multiple-testing correction.

### The broader peripheral signature is distinct from this neurological control cohort

A related question is whether the activation-independent PD signature above is PD-specific or simply reflects any peripheral neurological insult; we tested this genome-wide, across every cell type, using an independent neurological (not merely healthy) comparator, and revisit the same specificity question later at the resolution of a single Tem-CD8 subcluster.

Neurological Control patients are not a neutral, disease-free control, and not a homogeneous one either: 6 of the 9 had ischaemic stroke (blood drawn a median of 6 days post-diagnosis, range 3– 100 days), and the remaining 3 had other acute or subacute neurological presentations (C6 radiculopathy, myelitis, multifactorial gait disorder; Methods). Ischaemic stroke induces a distinct immune response (lymphopenia, compensatory suppression) mechanistically different from PD’s proposed chronic dysregulation, though this applies most directly to the stroke subset; these patients also had uncontrolled anticoagulant/steroid/antibiotic exposure, any of which could independently blunt a result in this group. So “absent in Neurological Control” should be read as evidence against a generic acute-CNS-injury explanation, not as a clean disease-free comparison (see Discussion).

**Supplementary Figure 7:**
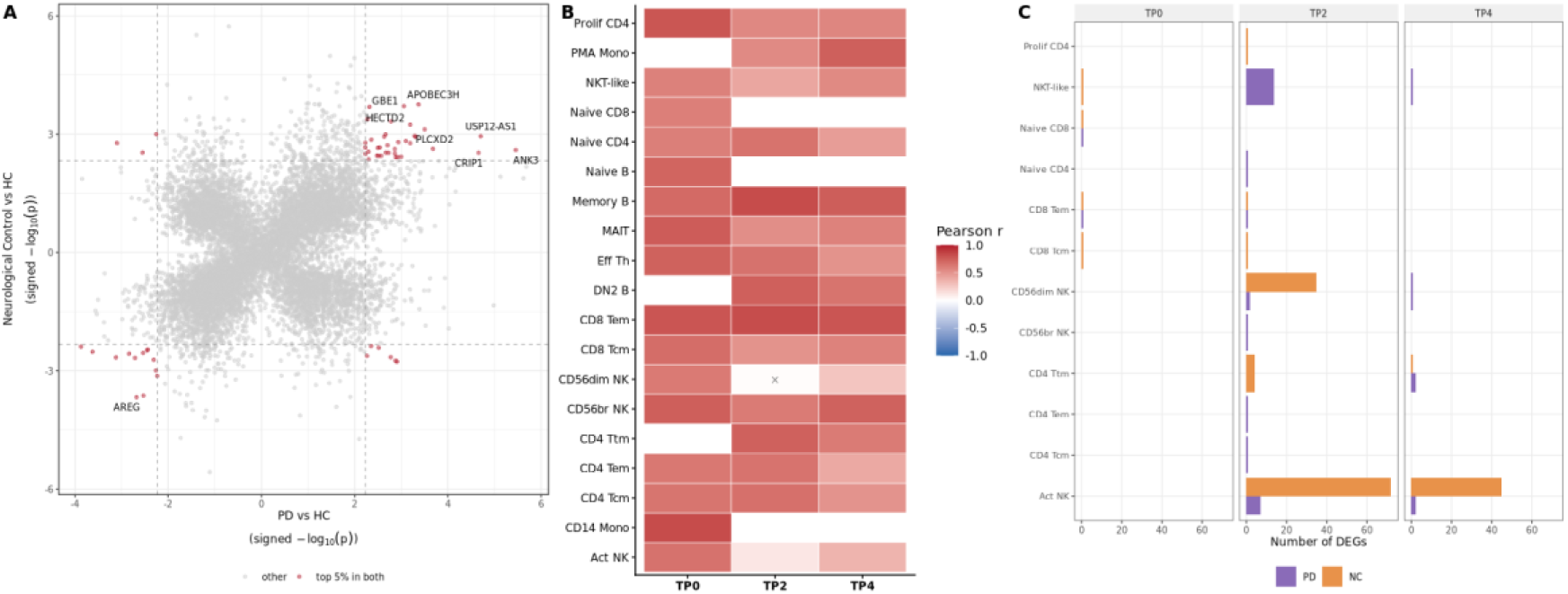
The broader peripheral signature is distinct from this neurological control cohort. (A) Threshold-free rank-rank overlap at TP4: every gene ranked by signed significance (-log₁₀ p x sign(log2FC), most extreme cell type per gene), independently for PD and Neurological Control vs. HC. Dashed lines mark each axis’s own top 5% cutoff (by |rank score|, universe = 10012 genes tested in both comparisons; n = 501 genes per top-5% list); red = in the top 5% of both rankings (n = 59, vs. 25.1 expected by chance; hypergeometric p <0.001)(B) Cross-disease LFC concordance heatmap (Pearson correlation, two-sided; × = BH adj. p ≥ 0.05). (C) DEG count per cell type and timepoint: PD vs. HC (red, n=14 PD, n=5 HC) and NC vs. HC (purple, n=9 NC, n=5 HC); limma-voom pseudobulk moderated t-test, two-sided, BH-adjusted. (see Methods, “Statistical power”).

The cross-disease comparison shows a specific signature, not merely a “nuanced pattern”: PD and neurological control changes are substantially correlated in overall direction and magnitude (median r = 0.69/0.64/0.58 at TP0/TP2/TP4), consistent with a shared inflammatory response upon stimulation, but diverge sharply in which individual genes reach significance (Supplementary Figure 7). At TP4, only 0 genes were shared between the PD and neurological control DEG sets, versus 6 PD-specific and 46 neurological control-specific genes; neurological control’s response is broader than PD’s at every timepoint (2/28/6 for PD vs. 4/114/46 for NC) and peaks at TP2 rather than TP4, concentrated in NK-lineage subtypes. This asymmetry may reflect a genuinely larger acute-injury response in the stroke-predominant Neurological Control group (Chamorro et al. 2012), or simply lower statistical power in this comparison rather than a true difference in effect size; either way, these counts are sensitive to the underlying FDR/fold-change threshold (Methods, “Statistical power”) and should be read as indicative of relative magnitude, not precise estimates.

With no genes shared at TP4, shared-gene enrichment could not be computed and the 6 PD-specific DEGs were too few for a well-powered ORA result, though the 46 neurological control-specific DEGs were enriched for retrotransposon silencing. Because this zero-overlap is itself a product of unequal power rather than necessarily a biological difference, we repeated the comparison threshold-free: ranking every gene by signed significance, the top 5% most-perturbed genes in each disease overlap far more than chance (Supplementary Figure 7 A; 59 shared vs. 25.1 expected, hypergeometric p <0.001; 50 concordant in direction), reconciling the strong LFC concordance (Supplementary Figure 7 B) with the earlier zero-overlap count (Supplementary Figure 7 C): the two conditions perturb an overlapping gene set, just not enough to clear a fixed significance bar in both cohorts given their very different sample sizes. Against this shared backbone, the activation-independent IKZF2 elevation in NK/NKT cells is disease-specific: it is absent in neurological control, and the same specificity check recurs later at the resolution of a single Tem-CD8 subcluster (a cleaner, patient-level test), with the same result.

## Stimulation Reveals a Cell-Type-Specific Failure to Resolve Activation

The sparse resting signal established above raises a sharper question: what happens to PD immune cells when they are actually challenged? PMA/ionomycin stimulation lets us ask this across every PBMC subset at once, forcing all major lineages through the same activation program simultaneously and following the response over the full time course.

### AP-1 induction is intact, but its late-window resolution fails in a specific subset of cell types

AP-1 (FOS/JUN dimers with EGR1–3, NR4A1–3, ATF3, and DUSP phosphatases, see “Methods: Module scores”) is the canonical transcriptional response to stress and antigen receptor signalling, and the gene family implicated in the Tem-CD8 exhaustion programme examined later. Its PD upregulation has been reported for monocytes and T cells individually; a recent resting-state atlas independently found an overlapping “activation signature” module elevated across most PD PBMC lineages at rest, including monocytes, CD56-bright NK, Tem-CD8, and MAIT cells (Moquin-Beaudry et al. 2025), confirming rather than newly reporting this elevation in resting Tem-CD8. We quantified AP-1 module scores across all PBMC subsets at each timepoint, applied a paired early/late-window test to every cell type, and examined sex stratification of the response.

**Supplementary Figure 8:**
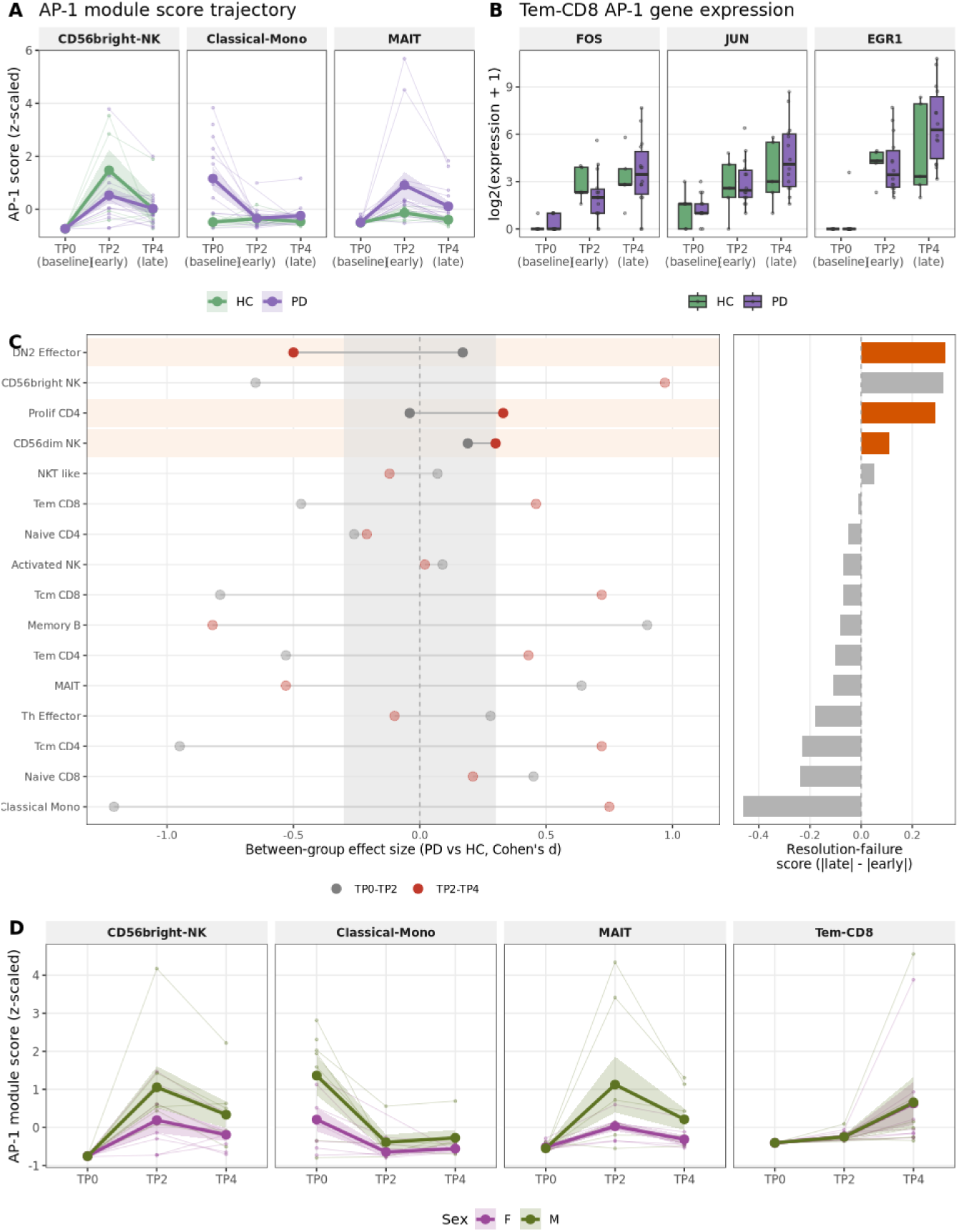
AP-1 pathway activation across PBMC subsets, its resolution failure over the activation time course, and sex stratification. (A) AP-1/EGR1 module score (mean expression of 17 AP-1 pathway genes, z-scaled within cell type) in HC (n=5) and PD (n=14) at TP0, TP2, and TP4, shown for three representative cell types spanning the lineages this section’s own prose identifies together (activated/Classical monocytes, CD56bright NK, MAIT) rather than every profiled cell type. Bold line + ribbon = group mean ± SE; faint lines = individual patients. (B) Raw expression of the same three representative AP-1 genes (FOS, JUN, EGR1) in Tem-CD8, the fourth curated cell type, shown separately as log2(expression + 1) boxplots by group and timepoint rather than as a z-scaled module score, since Tem-CD8 itself is examined in detail below. (C) Between-group (PD vs HC) AP-1 effect size in the early (TP0-TP2) and late (TP2-TP4) activation windows, across every cell type with sufficient donor pairs (n >= 3 per group). Cell types are ranked top-to-bottom by resolution-failure score (|between-group d| in the late window minus the early window), shown directly as a bar in the right-hand companion panel. Highlighted rows (left panel) are cell types where the early-window effect is itself small (|d| < 0.3, a genuinely comparable early response, grey reference band) and the late-window effect is at least moderate (|d| >= 0.3, outside that band); p is reported in Supplementary Table 3 for reference. (D) AP-1 module scores by sex and timepoint in PD patients, for the same four cell types as panels A and B; faint lines show individual patients behind the bold group mean ± SE.

AP-1 module scores were consistently elevated in PD relative to HC across all five major PBMC lineages (Supplementary Figure 8 A), a system-wide signal. The trajectory varied by cell type: monocytes were already elevated at TP0 in PD but converged with HC after stimulation, while Tem-CD8 diverged progressively through TP4 rather than plateauing. To test whether this is a Tem-CD8 idiosyncrasy or a general feature, we applied the same logic to every cell type with sufficient donor pairs (Supplementary Figure 8 C; Supplementary Table 3): a comparable paired rise from TP0 to TP2, diverging in the TP2-to-TP4 window. Ranking cell types by resolution-failure score (late-minus early-window between-group effect size) identifies 3 of 16 testable cell types where the early-window effect is itself small (|d| < 0.3, a genuinely comparable early response) and the late-window effect is at least moderate (|d| ≥ 0.3): DN2 Effector, Prolif CD4, CD56dim NK. Tem-CD8 does not itself meet this threshold in this genome-wide, AP-1-only sweep (resolution-failure score -0.01), but shows the same qualitative pattern far more clearly in the dedicated, Tem-CD8-specific paired test using its own composite AP1/exhaustion score (Supplementary Table 4, examined in detail below), this broader sweep establishes that unresolved, late-window-specific divergence is not unique to Tem-CD8, rather than being direct evidence for Tem-CD8 itself. At the individual-gene level, an unpaired, single-timepoint test (limma-voom, PD vs. HC) reached significance for none of them, at any of the three timepoints (minimum BH-adjusted p = 0.135) - underscoring why the paired, within-patient design above, not single-timepoint unpaired testing, is what recovers a real signal for this gene family.

AP-1 activation at TP4 follows the same sex-dominant variance structure established genome-wide later. Male PD patients show higher module scores in NK cells at TP4 (male = 360.07, female = 238.36; Wilcoxon p 0.209), the largest difference among the five lineages tested (Supplementary Figure 8 D); not apparent at rest (TP0: male = 10.84, female = 8.16), consistent with sex modifying response magnitude rather than baseline.

### Inferred cell-cell communication shows a sustainability failure across four functional axes at TP4

The AP-1 resolution failure described above is cell-intrinsic; we used CellChat ligand-receptor inference to test whether it is mirrored by a failure to sustain intercellular communication. Each condition/timepoint is a single pooled network, not per-patient replicates, so probability differences below describe the two pooled networks directly, not a between-patient comparison.

**Supplementary Figure 9:**
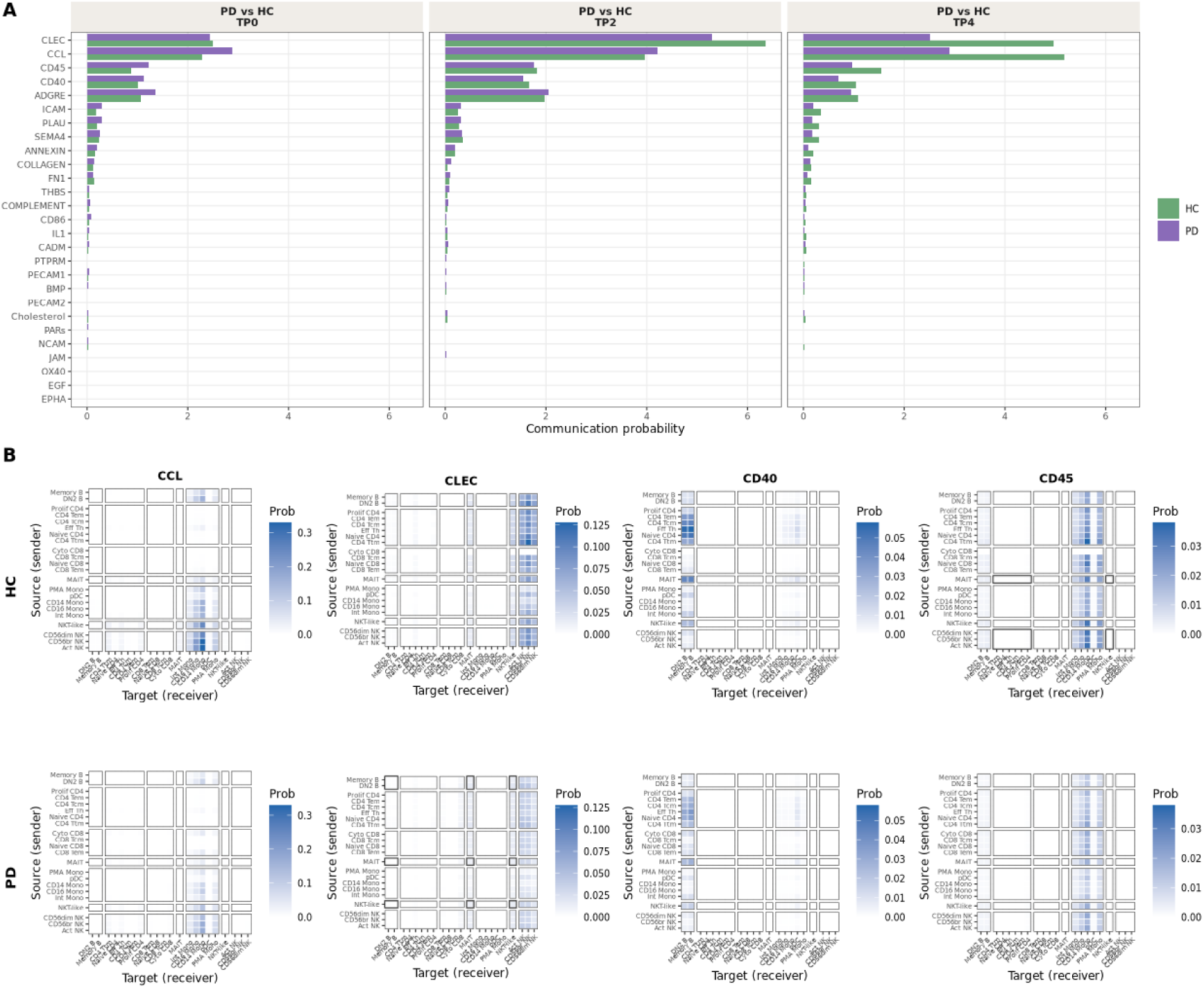
CellChat inferred communication analysis summary. (A) Absolute communication probability per pathway (PD vs. HC) at TP0, TP2, and TP4; pathways ordered by mean absolute HC-PD difference. (B) Sender-to-receiver communication probability heatmaps at TP4 for the four functional axes highlighted in detail (CCL, CLEC, CD40, CD45; columns, labelled), restricted to ligand-receptor pairs meeting CellChat’s own internal permutation-test threshold (p < 0.05) for being active within each condition’s pooled network, an internal test of pair activity, not a between-group statistical comparison.

Communication increased with stimulation in both groups, strongest in CCL, CLEC, ADGRE, CD45, and CD40 (Supplementary Figure 9 A); the early response was comparable (TP0: 0.401 PD vs. 0.337 HC; TP2: 0.618 vs. 0.641), but by TP4 PD was broadly reduced relative to HC (0.337 vs. 0.581).

This is a sustainability failure, not a blunted initial response: CCL and CLEC both failed to sustain their TP2-TP4 escalation in PD, reverting toward TP0 levels (CCL TP0: 2.89, TP4: 2.9 vs. HC 5.18; CLEC TP0: 2.43, TP4: 2.51 vs. HC 4.96). At the sender-receiver level for all four highlighted axes (Supplementary Figure 9 B), including CD40 (PD 0.7 vs. HC 1.04) and CD45 (PD 0.97 vs. HC 1.55): all four show a lower absolute communication probability in PD than HC specifically at TP4. This is not explained by cell counts (PD ∼36,000 cells at TP4 vs. HC’s ∼13,400). This rules out a simple cell-number shortfall, but cell-type-proportional differences between groups were not fully controlled for here and may still contribute to the apparent pathway-level differences.

PD immune cells therefore activate normally by TP2 but fail to sustain predicted signalling through TP4, a functional correlate of the transcriptional resolution failure above.

## Convergence: Sex-divergent Signals Meet in Tem-CD8

The activation-independent PD signature established earlier in this manuscript is broad and holds up against an independent neurological comparator (above); under stimulation, PD immune cells mount a normal initial response but a specific subset of cell types fails to sustain it, with parallel consequences for inferred intercellular communication (above). One variable cuts across all of this and has received surprisingly little attention in PD immunology: sex, and how it shapes the variance structure of PD PBMCs and the trajectories male and female patients follow as cells respond to stimulation, down to individual genes rather than composite scores alone.

### The variance structure of PD PBMCs is dominated by sex, not disease severity

We decomposed transcriptional variance within PD at each timepoint using PVCA (Methods), now restricted to PD alone so that sex and disease severity can be compared directly. At rest (TP0), the two largest non-residual sources of variance were Sex (35.1%) and clinical_type_description:sex (22.7%). Within PD, sex remains the dominant source of transcriptional variance at every individual timepoint (TP0 = 35.1%, TP2 = 45.6%, TP4 = 43.5%), while Hoehn-Yahr Stadium alone explains less than 1% at any timepoint (Figure 6 A) and its interaction with sex adds little more (2.2%/36.7%/38.7%; combined with the sex main effect: 37.3%/82.3%/82.2%).

**Figure 6:**
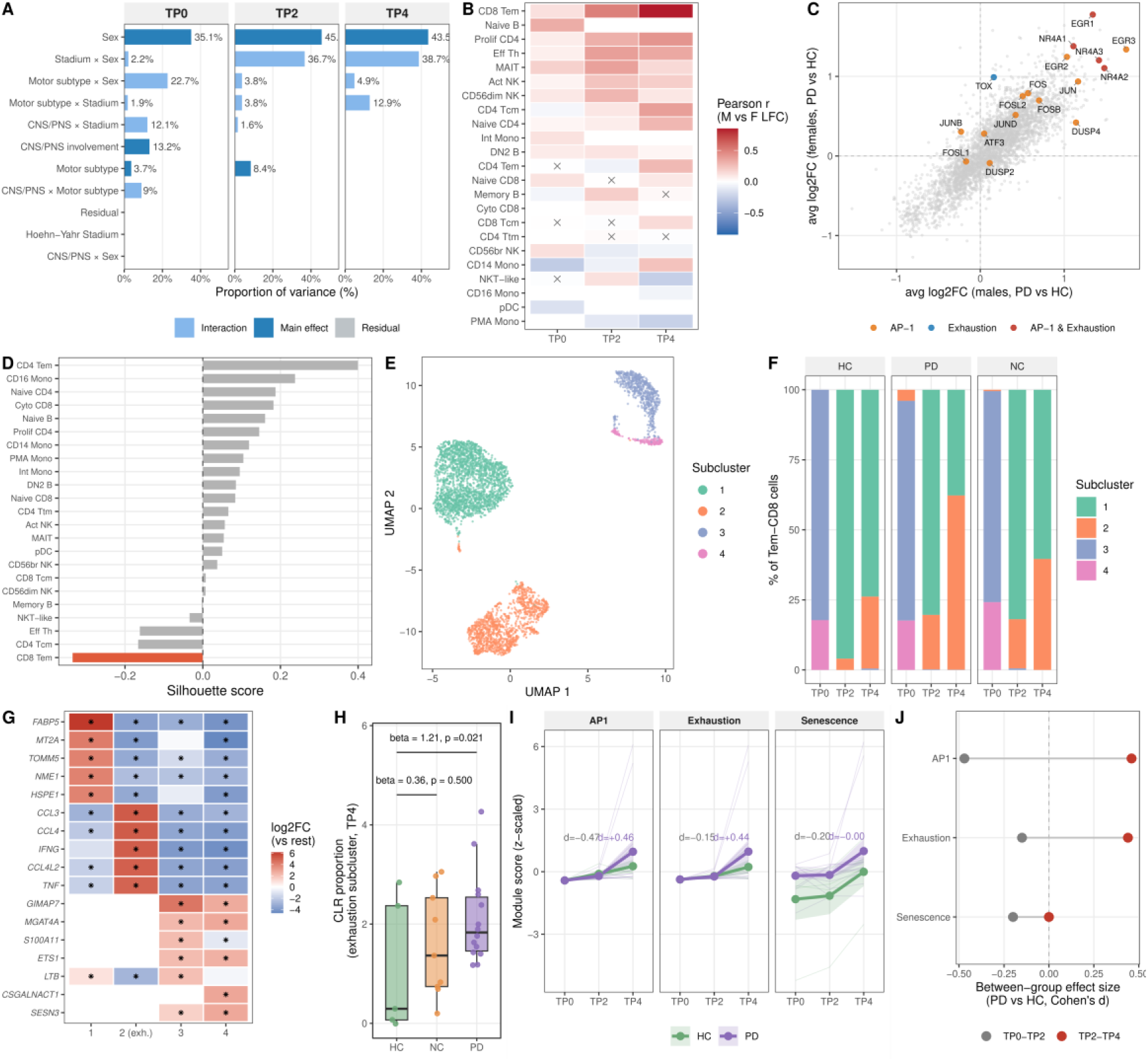
Sex-stratified analysis of the Tem-CD8 population across the activation time course (PD: n=7 male, n=7 female; HC: n=3 male, n=2 female; panel F additionally includes Neurological Control, n=S). (A) Within-PD PVCA by timepoint (pvcaBatchAssess threshold 0.75, per-timepoint model), quantifying variance attributable to sex versus Hoehn-Yahr Stadium. (B) Cross-sex LFC concordance (two-sided Pearson correlation, r, on male vs. female Wilcoxon rank-sum avg_log2FC, PD vs. HC; × = BH-adjusted p ≥ 0.05) by cell type and timepoint. (C) Male vs female Tem-CD8 log2FC (PD vs HC) at TP4; AP-1 and exhaustion gene sets highlighted. (D) Silhouette scores per cell type. (E) Tem-CD8 re-clustered on its own variance structure (4 subclusters; uncorrected PCA, not timepoint-corrected - see Methods). (F) Subcluster frequency by group and timepoint: subcluster 2 (elevated Exhaustion/AP1 module scores; top markers CCL3, CCL4, IFNG, CCL4L2, TNF, NR4A3) makes up c2.2% of PD Tem-CD8 cells at TP4 versus 25.7% in HC and 3S.c% in Neurological Control. (G) Top marker genes per subcluster (one-vs-rest pseudobulk limma-voom; * = BH adj. p < 0.05). (H) Differential abundance of the exhaustion subcluster at TP4: per-patient CLR-transformed proportions (age-adjusted linear model) - the patient-level, pseudoreplication-free measure of the compositional shift. (I) AP1, Exhaustion, and Senescence module score trajectories in Tem-CD8 (mean +/- SE per group; faint lines = individual patients); annotated d = between-group (PD vs HC) effect size (Cohen’s d) on the within-patient change in that window (grey = early window TP0-TP2, red = late window TP2-TP4; patient-level; full stats in Supplementary Table 4). (J) The same between-group effect sizes for all three scores shown directly, ranked by score rather than cell type (companion to panel I and Supplementary Table 4; grey = early window TP0-TP2, red = late window TP2-TP4).

### Sex-stratified disease signals are concordant in direction

Cross-sex LFC concordance (Methods) quantifies whether male and female patients’ PD-vs-HC fold-changes point the same way, gene by gene, a descriptive statistic, not a significance test, given the small size of each sex-stratified stratum.

At baseline (TP0), cross-sex concordance of PD-associated gene expression changes was near zero across virtually all cell types (Figure 6 B), indicating a largely sex-dimorphic transcriptional response to PD at rest. Two cell types illustrate the qualitative range: Classical Mono showed the strongest anti-concordance (r = -0.27: male and female PD patients dysregulate the same genes in opposite directions), while Naive B showed the highest, though still weak, positive concordance at TP0 (r = 0.33).

Concordance increased over time in several cell types, most strikingly in Tem-CD8 (r = 0.12 → 0.51 → 0.85, reaching near-complete convergence by TP4), with more modest increases in Prolif-CD4 (0.07 → 0.31 → 0.42) and Tcm-CD4 (−0.04 → 0.1 → 0.38). A permutation test (shuffling gene labels in one sex’s LFC vector 500 times, G-04) confirmed the TP4 concordance (r = 0.85) exceeds chance: the observed r exceeded the 95th percentile of the null distribution (0.031, permutation p = 0).

The concordance analysis also flagged a Tem-CD8-specific signal at TP4: 340 genes were nominally significant in both sexes independently in the wilcox analysis (100% directional concordance, LFC Pearson r = 0.958) yet undetectable in limma-voom pseudobulk; wilcox p-values alone are pseudoreplication-prone, but given the already-high overall cross-sex correlation at TP4 (above), this near-total directional agreement is at least consistent with a real signal rather than independent shared false positives. The top concordant genes (NR4A1, NR4A2, NR4A3, EGR1, EGR3) are canonical AP-1/exhaustion transcription factors, a biologically coherent set suggesting a real signal diluted below pseudobulk detection when averaged across the full Tem-CD8 pool (Figure 6 C). Combined with the high cross-sex concordance at TP4 (r = 0.85), this was a preliminary indication motivating the focused subclustering analysis below, to test whether a discrete Tem-CD8 subpopulation underlies it.

Further supporting this, the exhaustion/AP-1 transcription factor set itself (TOX, TOX2, NR4A1/2, PDCD1, TIGIT, LAG3, FOS, FOSB, JUN, JUNB, BATF, EGR1) shows near-identical mean log2FC in Tem-CD8 by TP4 between male PD (1.16) and female PD (1.72).

This rise in cross-sex concordance is not confined to Tem-CD8, Prolif-CD4, and Tcm-CD4: of 16 cell types with concordance data at all three timepoints, 11 show higher concordance at TP4 than at rest (mean r 0.04 at TP0 vs. 0.2 at TP4), a broad convergence toward a shared programme by peak activation, with one clear exception: an innate/myeloid axis that remains or becomes sex-discordant at TP4 (NKT like, PMA Mono, CD56bright NK, NonClassical Mono, Ttm CD4, 5 cell types in total, all negative). This convergence reflects parallel rather than shared upstream trajectories, much as male and female PD patients present with different symptom profiles and progression despite a shared diagnosis (Cerri et al. 2019); Tem-CD8’s exhaustion/AP-1 programme is simply its most complete instance.

### The exhausted state deepens through activation

To test whether the Tem-CD8 exhaustion signal is a positive, graded phenomenon rather than a DEG-analysis artefact, we computed a per-cell exhaustion module score (AddModuleScore over NR4A1, NR4A2, NR4A3, TOX, LAG3, TIGIT, PDCD1, HAVCR2, EGR1) across 4121 Tem-CD8 cells from 28 patients. The PD-HC gap widens across the time course (0.1/0.15/0.58 at TP0/TP2/TP4) and at TP4 is 2.8× the equivalent Neurological-Control-vs-HC gap (0.21), an early indication of disease specificity. A per-cell Wilcoxon test reaches significance at every timepoint (p <0.001/<0.001/<0.001), though such tests can overstate confidence by treating within-patient-correlated cells as independent. Consistent with a real effect concentrated in a subpopulation of Tem-CD8 cells rather than a uniform shift across the whole population, the signal is no longer significant once patients are the unit of analysis (patient-level p 0.785/0.500 at TP2/TP4; 2 of 14 PD patients exceed the highest-scoring control at TP4). To isolate that subpopulation directly, we turn to subclustering below.

### Tem CD8 subclustering

Cluster separation quality, assessed by silhouette scoring across all annotated cell types, independently motivated the subclustering analysis: Tem-CD8 had the lowest silhouette score of any cell type (score = -0.34; Figure 6 D), indicating these cells sit, on average, closer to a neighbouring cluster than their own centroid, a quantitative signature of internal heterogeneity. Tcm-CD4 and Th-Effector had slightly negative silhouette scores too (−0.17 and -0.16), but neither shows the cross-sex convergence signal that specifically motivated Tem-CD8 (r = 0.38 and 0.34 vs. 0.85 at TP4), so Tem-CD8 alone was re-clustered.

The subclustering resolves this directly (Figure 6 E-F): Tem-CD8 cells split into 4 subclusters when re-clustered on their own variance structure (Methods). One subcluster stands out sharply on both Exhaustion and AP1 module scores (1.07 and 0.88, vs. near-zero or negative in the others). It combines a canonical checkpoint/exhaustion transcription factor signature with heightened, not silenced, effector cytokine and chemokine output: a mixed picture, not the classical hyporesponsiveness that exhaustion implies in its strict sense (Discussion). Its top marker genes (CCL3, CCL4, IFNG, CCL4L2, TNF, NR4A3; Figure 6 G) span the NR4A/immediate-early family alongside CCL3/CCL4/IFNG/TNF. These same effector cytokines’ inferred cross-cell communication fails to sustain itself specifically in PD at TP4 (Supplementary Figure 9).

This subcluster is where the sex-concordant, PD-specific exhaustion signal anatomically lives: by TP4 it comprises 62.2% of PD Tem-CD8 cells vs. 25.7% in HC and 39.6% in Neurological Control. This is confirmed by a differential abundance test on CLR-transformed, per-patient proportions (age-adjusted, 28 patients, the same method used elsewhere in this manuscript): PD differs significantly from HC (β = 1.21 on the CLR scale, p 0.021) while Neurological Control does not (β = 0.36, p 0.500) (Figure 6 H). Since mean module scores within the subcluster are similar across groups (PD 1.1, HC 1.01, Neurological Control 1.02), this indicates that PD shows more cells that become this subcluster rather than making its cells more exhausted. This supports the hypothesis above: the sparse baseline DEG landscape described earlier partly reflects analysing a heterogeneous cell pool whose disease-relevant subpopulation only declares itself under stimulation.

### The paired activation trajectory explains why this subcluster accumulates in PD

Why this subcluster accumulates specifically in PD is explained by a within-patient paired analysis of the same AP1 and Exhaustion module scores across the two activation windows (Figure 6 I; Supplementary Table 4), the same early-parity, late-divergence pattern already established for AP-1 more broadly (Supplementary Figure 8 C; Supplementary Table 3), now shown to hold in the population and subpopulation where it culminates. From TP0 to TP2, PD and HC show comparably large paired increases in both exhaustion (dz = 0.91 vs. 1.33) and AP1 score (dz = 1.22 vs. 1.04), with no between-group difference (d = -0.15 and -0.47). From TP2 to TP4 the groups separate: PD continues to escalate (exhaustion dz = 0.62, p <0.001; AP1 dz = 0.61, p <0.001) while HC’s change is smaller and directionally inconsistent (exhaustion dz = 0.53, p 0.812; AP1 dz = 0.41, p 0.812), producing a moderate divergence confined to this window (d = 0.44 and 0.46). The cumulative TP0-to-TP4 gap (d = 0.4 for exhaustion, 0.38 for AP1) is therefore driven almost entirely by this late window, not a uniformly larger PD response from the start, the same pattern re-emerges genome-wide below once power is recovered by pooling HC with Neurological Control.

A parallel test on a CD8 T cell senescence module score (Methods, “Module scores”) shows that early-window changes are comparable between groups (dz = 0.06 PD vs. 0.48 HC; d = -0.2), and late-window changes are dz = 0.61 PD vs. 1.65 HC (d = 0): this does not reproduce the AP-1/exhaustion pattern (diverging in the early window as well, or in the opposite direction). This does not support treating senescence as empirically established in this population to the same extent as exhaustion.

### The composite-score divergence generalises genome-wide

Supplementary Table 4 established this divergence using two hand-built composite scores; to test whether it holds gene-by-gene rather than as an artefact of gene-list curation, we repeated the same paired difference-of-coefficients test genome-wide, pooling HC with Neurological Control (n=14, inverse-variance meta-analysis; Methods) to recover the power HC alone (n=5) lacks, a check on an already-established signal, not independent replication, since it assumes the two groups’ paired trajectories are homogeneous enough to combine (only partly testable; Supplementary Figure 10 B).

**Supplementary Figure 10:**
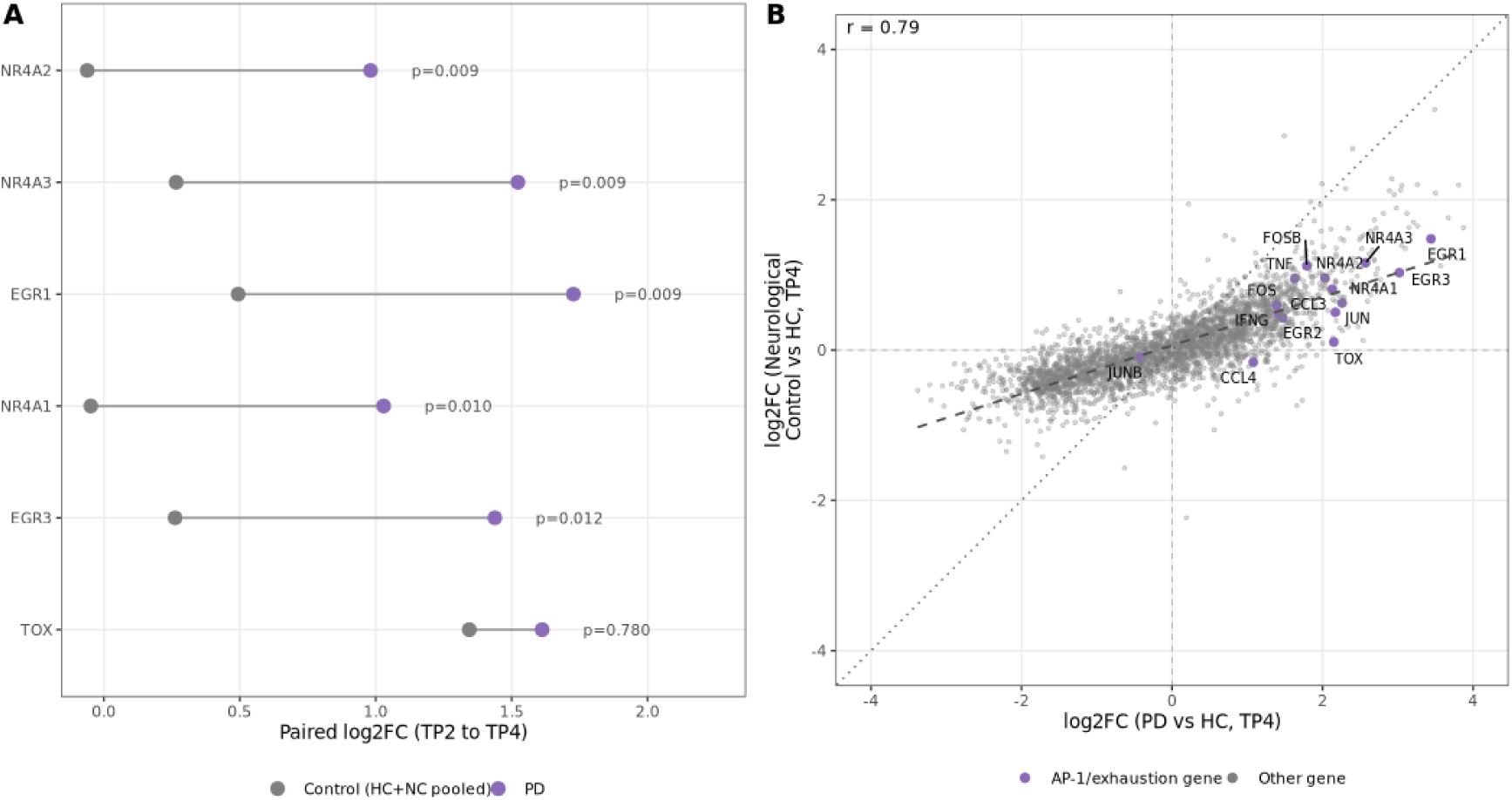
Genome-wide validation of the Tem-CD8 TP2-to-TP4 divergence using pooled controls. (A) Gene-level paired log2FC (TP2-to-TP4 within-patient change) for the pooled control group (HC + NC, n=14, inverse-variance meta-combined) versus PD (n=14), from the pooled-control validation of the Tem-CD8 genome-wide divergence test (two-sided difference-of-coefficients test, Welch-Satterthwaite-approximated t-distribution; Methods, ‘Within-group activation trajectories’). (B) A check on the homogeneity assumption in (A) using data that actually distinguishes the two control sources (unlike the pooled export in A, which only retains their already-combined estimate): cross-sectional log2FC at TP4 (PD vs HC; NC vs HC) restricted to Tem-CD8, one point per gene (n = 3083; Pearson r = 0.79).

This recovers a significant TP2-to-TP4 divergence in 5 of 6 genes (Supplementary Figure 10 A; NR4A1, NR4A2, NR4A3, EGR1, EGR3; BH-adjusted p = 0.009-0.012), converging on the same canonical AP-1/exhaustion genes already implicated by the composite scores.

## Discussion

This study provides a time-resolved, sex-stratified single-cell map of ex vivo-activated PBMCs in Parkinson’s disease. Male and female PD patients show largely divergent transcriptional states at rest, and sex explains more of that variance than disease status across most cell types. This dimorphism largely resolves after activation, converging on a shared programme in CD8+ effector memory T cells with an exhaustion-associated transcriptional component with a co-occurring senescence signature tested directly but not supported to the same extent. The same late-window pattern recurs independently in AP-1 induction and in a coordinated drop in inferred intercellular communication at TP4, and is not observed in the neurological controls, consistent with a disease-rather than injury-associated effect. The PD peripheral immune phenotype is therefore better understood as a sex-stratified dynamic response than a fixed, resting-state signature.

The Tem-CD8 exhaustion-associated programme illustrates why prior studies missed this: it is undetectable in our own TP0 data, and it is absent from a resting-state PD PBMC study that found no CD8+ exhaustion signature, only an enhanced cytotoxic/differentiated phenotype in females at rest (Capelle et al. 2023). That finding sits uneasily alongside reports of reduced CD8+ senescence markers in early PD (Williams-Gray et al. 2018; Kouli et al. 2021). These are incomplete rather than contradictory: an activation-gated state cannot be detected in unstimulated blood, regardless of power. The closest precedent, an ex vivo study stimulating isolated T cells with CD3/CD28 and monocytes with IFN-γ, similarly found PD-specific deficits only after stimulation — lower mitochondrial health specifically in CD8+ T cells, and reduced cytokine output from bulk stimulated T cells (Mark et al. 2025) — but used a single post-stimulation timepoint, metabolic assays rather than transcriptomics, and no exhaustion markers. Antigen-experienced, clonally expanded CD8+ T cells have separately been found in the cerebrospinal fluid of both Alzheimer’s and, in a small cohort, Parkinson’s disease patients (Gate et al. 2020). This is consistent with growing evidence that such cells can occur in neurodegenerative disease more broadly, although whether they correspond to the exhaustion-associated state identified here remains unknown.

Even within our own time course, the signal was confined to a narrow window (Tem-CD8 divergence concentrated specifically in TP2-to-TP4, mirrored by the inferred communication drop at the same timepoint), so a single post-stimulation snapshot risks missing it entirely. AP-1 activation is not exclusively activation-revealed, however: an independent resting-state atlas found an overlapping activation-signature module already elevated in the same PD compartments (Moquin-Beaudry et al. 2025), confirming rather than originating the observation.

Disease severity follows the same principle, independent of sex: Hoehn-Yahr Stadium correlates with few genes at rest but far more at peak activation (1701 pairs at TP0 vs. 17911 at TP4), led by NR4A2 in central-memory CD4 T cells and HMOX1, a cytoprotective oxidative-stress/PD-risk gene. NR4A2’s positive correlation with Stadium at rest (r = 0.55) sits in tension with reports of reduced resting NURR1/NR4A2 in mixed peripheral lymphocytes (Le et al. 2008; Liu et al. 2012), plausibly reconciled by cell-type specificity (Tcm-CD4-specific here). Even healthy PBMCs follow this pattern at the whole-transcriptome level: an early, shared immediate-early wave gives way to a narrower, cell-type-specific late wave. A field still dominated by resting-state profiling may therefore be systematically underestimating dysfunction that only a time-resolved challenge reveals.

Sex is this dataset’s second organising axis, and one PD research has largely overlooked, despite males developing PD at roughly twice the rate of females and females progressing faster with a different response to dopaminergic therapy (Cerri et al. 2019). Existing PD PBMC transcriptomic studies have mostly been underpowered to stratify by sex, or treated it as a nuisance covariate (Xiong et al. 2024; Wang et al. 2021). Sex-dimorphic changes are reported in PD brain tissue (López-Cerdán et al. 2022; Tranchevent et al. 2023), but neither this degree of dimorphism nor a variance-partitioning comparison against disease status had previously been shown in peripheral blood. Our data close that gap: PVCA shows sex explains more transcriptional variance than disease status at every timepoint (35.1%/45.6%/43.5% vs. <1% for Stadium). Male and female PD patients also show largely divergent responses at rest, most strikingly in 5 cell types (NKT like, PMA Mono, CD56bright NK, NonClassical Mono, Ttm CD4; Classical Monocytes itself is r = 0.24 at TP4), with negative cross-sex concordance at TP4, a pattern that would cancel out entirely in a pooled, sex-uncontrolled analysis. The biological basis of this dimorphism remains unresolved: sex-dependent hormonal and pharmacological effects are plausible contributors, given known sex differences in immune regulation (Cerri et al. 2019) and in levodopa pharmacokinetics (women show higher plasma exposure at equivalent doses (Kumagai et al. 2014)), but cannot be distinguished in the present cohort. Against this backdrop of pervasive divergence, one signal resolves it entirely, and it is the most unexpected result in this study: Tem-CD8 responses, essentially uncorrelated between sexes at TP0, converge until male and female log2FC vectors are nearly collinear by TP4. Whether sex-dependent hormonal signalling contributes to this shared endpoint is an interesting question for future studies.

What does this shared endpoint actually represent? Part of it overlaps with the canonical exhaustion programme at the level of transcriptional regulators and checkpoint receptors: TOX is a central transcriptional and epigenetic regulator of the exhaustion programme in chronic infection and cancer (Khan et al. 2019), and the NR4A family (NR4A1/2/3, all three upregulated here) has been shown to promote this dysfunctional state by antagonising AP-1-dependent effector transcription (Chen et al. 2019; Liu et al. 2019). This provides a potential mechanistic connection to the unresolved AP-1 induction observed here. Together with checkpoint receptors (HAVCR2, LAG3, TIGIT), this follows a transcription factor hierarchy proposed for both chronic viral infection and tumour-infiltrating lymphocytes (Blank et al. 2019) — though whether a single hierarchy generalises cleanly across both contexts is itself debated — canonically triggered by persistent antigen exposure and sustained inflammatory signalling (Wherry and Kurachi 2015). But we did not find classically exhausted T cells: the subcluster’s top marker genes are effector cytokines and chemokines (CCL3, CCL4, IFNG, TNF): heightened output, not the cytokine-silenced signature exhaustion implies in its strict sense. This is expected given the stimulus: PMA/ionomycin bypasses the TCR entirely and drives cytokine transcription directly via PKC/calcium (Truneh et al. 1985), so an assay built to maximise effector output cannot, on cytokine readouts alone, distinguish exhaustion from hyperactivation. This is why “exhaustion-associated,” is used deliberately throughout this manuscript to describe a checkpoint-co-expressing, hyperactivated effector state rather than classical terminal dysfunction. Its PD-specificity may reflect prior in vivo antigen exposure poising these cells toward this state, consistent with α-synuclein-reactive T cells in PD blood (Sulzer et al. 2017), but the transcriptional data alone can’t resolve this; functional validation and TCR repertoire data are natural next steps.

Several observations argue against this being a non-specific consequence of strong stimulation rather than a PD-associated process: cross-sex concordance rises progressively rather than starting high (r = 0.12 → 0.51 → 0.85), TP4 concordance exceeds the 95th percentile of a gene-permutation null (0.031, permutation p = 0), and neurological control patients under the same protocol don’t show it. Differential survival (a pre-existing exhausted-like subset preferentially surviving 4-hour culture in PD) is not supported by a direct check: mitochondrial read fraction, a standard per-cell stress/death proxy, did not differ between the exhaustion subcluster and the rest of Tem-CD8 within PD patients at TP4 (paired by patient, Wilcoxon signed-rank p = 0.376, n = 13), nor between PD and HC within the exhaustion subcluster itself (p = 0.084). This cannot fully exclude differential survival, since standard QC filtering would already have removed any cell too far gone to be captured, but it argues against a coarse, detectable stress signature driving the effect. A related question is whether this state is activation-revealed or activation-induced: if merely unmasked, a patient’s resting-state score should predict their activated-state score. It did not: TP0 exhaustion module score (the same 9-gene panel as Supplementary Table 4) did not predict TP4 score within our paired design (Spearman rho = 0.08, p = 0.781, n = 14 PD; rho = 0.26, p = 0.668, n = 5 HC, too few to interpret alone). This argues against a straightforward quantitative carry-over from the resting-state exhaustion score, but does not distinguish induction from unmasking. A null result at this size can’t rule out a true signal too small to detect, and genuine validation needs an independent activation cohort, not a resting-state comparison.

The disease-specificity claim rests partly on the neurological control comparator, included to distinguish PD-specific immune alterations from generic CNS injury responses. The Tem-CD8 programme’s absence there supports disease association, corroborated genome-wide, but the comparator is heterogeneous and predominantly post-stroke (6 of 9 patients; Methods), whose acute DAMP-mediated immune response and post-ischaemic immunosuppression are mechanistically distinct from PD’s proposed chronic, antigen-driven dysregulation (Chamorro et al. 2012). So the signature’s absence here supports a chronic-neurodegeneration-associated interpretation rather than a generic acute-injury one, but cannot establish specificity against every alternative condition; comparison against other parkinsonian conditions (one PD PBMC atlas found its AP-1 signature absent in atypical parkinsonism (Moquin-Beaudry et al. 2025)) would strengthen this further.

Several further limitations qualify these conclusions. Cohort size is foremost: with 14 PD patients and 5 healthy controls, the study is underpowered for sex-stratified single-timepoint comparisons; paired within-patient designs were used wherever possible to mitigate this, but negative findings should still be interpreted cautiously given the limited power. Two additional factors warrant consideration for the sex-dimorphic signal specifically: all PD patients were medicated, and medication load correlated with disease severity (Spearman ρ = 0.8, p = < 0.001). Dopaminergic drugs have been proposed to modulate T cell function (Baird et al. 2019), and since dopamine receptor expression is itself sex-specific, medication could plausibly mimic or amplify part of the signal reported here. HLA genotype was computationally inferred from the scRNA-seq reads but could not be evaluated as a covariate at this cohort size. This remains relevant since HLA variants, including HLA-DRB5, are among the loci associated with PD risk (Nalls et al. 2019) — HLA molecules more generally shape antigen presentation — and α-synuclein-reactive T cells in PD are MHC-restricted (Sulzer et al. 2017); larger cohorts integrating HLA genotype with TCR specificity are a natural next step.

The inferred cell-cell communication analysis carries its own caveats: CellChat’s predicted TP4 reduction in ligand-receptor communication probability (CCL, CLEC, CD40, CD45) reflects expression data, not measured secretion or receptor occupancy. Each condition/timepoint is also a single pooled network without per-patient replicates, so no between-patient variance estimate exists. PMA/ionomycin itself measures maximal intracellular activation potential downstream of PKC and calcium signalling, not immune competence broadly, and does not extrapolate to physiological antigen-specific responses. PKC isoforms and calcium-signalling components are themselves lineage-restricted (Meller et al. 1998; Isakov and Altman 2002; Feske 2007), and memory/effector T cells carry a pre-poised chromatin state naive T cells lack (Northrop et al. 2006). This explains the muted, high-variance monocyte response relative to T/NK/MAIT cells: lineage-specific pathway engagement, not a technical failure. It is also consistent with the shared emergence of a distinct “PMA-induced monocyte” population across all three groups by TP4, an expected consequence of strong stimulation rather than a PD-specific effect.

Disease-stage heterogeneity is a related concern: PD patients spanned a range of Hoehn-Yahr stages, whether the Tem-CD8 programme itself tracks severity wasn’t formally tested, and this cross-sectional design can’t separate stage from disease duration. Transcriptomics is also only a proxy for function here: the exhaustion label reflects transcriptional similarity to a canonical gene set, not a functional measurement.

Finally, cryopreservation freeze duration differed between the PD and HC cohorts but was not associated with cell recovery or transcriptional profile at any timepoint (Results), ruling it out as a technical confound within our own cohort. Separately, no prior study used this exact activation time-course design, so external replication is necessarily indirect, and comparability is limited across every available comparator: differently-stimulated cohorts (an ex vivo CD3/CD28-stimulated CD4+ T cell bulk RNA-seq time-course (Diener et al. 2023); a CD3/CD28-and-IFN-γ-stimulated study using metabolic rather than transcriptomic assays (Mark et al. 2025)) and resting-state PBMC datasets (Grandke et al. 2025; Capelle et al. 2023; Moquin-Beaudry et al. 2025) all differ from our own protocol in stimulus, cell type, or activation state.

Replication in a prospective, pre-registered cohort is the priority next step, ideally paired with protein-level validation of the checkpoint/effector co-expression pattern and TCR repertoire data to test whether this state tracks antigen-experienced, clonally expanded T cells. Whether the Tem-CD8 programme is detectable in an independent tissue (e.g. postmortem brain-resident T cells), and whether the exploratory peripheral-to-brain ligand-receptor bridging reflects a real interaction, also remain untested.

Taken together, this study’s throughline is not that PD patients carry an unusual T cell population, but that their immune system responds differently once challenged. That difference is visible only in the dynamics of the response: healthy PBMCs mount and resolve an activation programme, PD monocytes partially normalise, and PD Tem-CD8 cells instead fail to resolve toward the same late-state trajectory, arriving instead at a sex-convergent, exhaustion-associated state. Resting blood, still the dominant paradigm in peripheral immune phenotyping of PD, may therefore systematically underestimate dysfunction that only becomes visible under challenge. The Tem-CD8 state provides a candidate target for mechanistic investigation and, with independent validation, future biomarker development. More broadly, longitudinal activation profiling, sex stratification, and appropriate inflammatory comparators may provide a useful framework for studying peripheral immune dysfunction in other neurodegenerative and neuroinflammatory conditions.

## Methods

### Study cohort and ethics

Participants were recruited at Saarland University Medical Center (Universitätsklinikum des Saarlandes, UKS); the pilot cohort was recruited in April–May 2022, and the main cohort from May 2022 through January 2023. PD patients were clinically diagnosed with idiopathic Parkinson’s disease by their treating neurologist per routine clinical judgment. The donors (donors 1–4) of the initial pilot analyses were a group of female volunteers, all 22 years of age. PD patients (n=14) were elderly persons at different stages of the disease (equivalent rates of male and female; mean 66 years; +15/−22 years). The control cohort was matched for age (mean 72 years; +15/−16 years) and sex and comprised healthy volunteers (n=5) and Neurological Control donors (n=9) recruited for acute or subacute neurological conditions: ischaemic stroke (n=6), C6 radiculopathy (n=1), myelitis (n=1), and multifactorial gait disorder (n=1). Among the stroke patients, blood was drawn a median of 6 days after diagnosis (range 3–100 days). Basic clinical data for all participants, including PD manifestation, Hoehn-Yahr stage, and disease duration, are summarised in Supplementary Table 1 (Metadata); comorbidity-relevant concomitant medication is detailed in the full per-patient medication panel (Supplementary Figure 6); no patient was on an immunosuppressant medication. Peripheral blood was drawn by venepuncture into lithium heparin-containing collection tubes (S-Monovette; Sarstedt AG & Co. KG, Nümbrecht, Germany); a total of 27 ml of peripheral blood was collected per subject. The study was approved by the Research Ethics Committee of Saarland University (ethical vote number: 48/22). Written informed consent was obtained from all participants.

This was an observational, non-interventional cohort study: participants were not assigned to study arms or treatment conditions, so randomization was not applicable. Sample processing and sequencing were not performed blind to disease/control status: sample identifiers made group identity apparent to whoever handled each sample at every step. Samples from PD, HC, and NC donors were processed in mixed order rather than batched by group.

All PD patients were on dopaminergic medication at the time of sampling (14/14; Supplementary Figure 6); medication load was tested for correlation with Hoehn-Yahr Stadium (Spearman). The potential confounding effects of medication and HLA genotype on the transcriptional findings are discussed in the Discussion.

### PBMC isolation and cryopreservation

PBMCs were isolated from peripheral blood by density gradient centrifugation using Pancoll human density gradient medium (ρ = 1.077 g/ml; PAN-Biotech GmbH, Aidenbach, Germany). Isolated cells were resuspended in 1× Dulbecco’s Phosphate Buffered Saline (DPBS; Gibco™, Thermo Fisher Scientific Inc., Waltham, MA, USA) and the total number of viable cells was determined by trypan blue staining (Gibco™ Trypan Blue Solution, 0.4%; Thermo Fisher Scientific Inc.) and haemocytometer brightfield counting (Incyto X50, Improved Neubauer; Thermo Fisher Scientific Inc.).

For cryopreservation, approximately 5 × 10^6^ viable cells per aliquot were pelleted and resuspended in a 1:1 (v/v) combination of cryopreservation mixes A and B. Cryopreservation mix A consisted of 50% (v/v) RPMI 1640 (Gibco™, Thermo Fisher Scientific Inc.) and 10% (v/v) heat-inactivated fetal bovine serum (FBS; Biochrom GmbH, Berlin, Germany). Cryopreservation mix B contained 80% (v/v) heat-inactivated FBS and 20% (v/v) dimethyl sulfoxide (DMSO; Carl Roth GmbH & Co. KG, Karlsruhe, Germany). Cells were frozen at −80°C for 1–3 days in an isopropanol-containing cryo-freezing container and subsequently transferred to a liquid nitrogen tank for long-term storage. Cellular samples used for single-cell sequencing analyses were cryopreserved for less than 9.5 months.

### PMA-Ionomycin stimulation and methanol fixation

Cryopreserved cell stocks were thawed at 37°C in a water bath and pelleted (1,600 rpm, 6 min) to replace the cryopreservation mix with 500 µl of pre-warmed RPMI 1640 medium supplemented with 10% (v/v) heat-inactivated FBS and 1% (v/v) penicillin-streptomycin (Gibco™, 100 U/ml; Thermo Fisher Scientific Inc.). The cell suspension was transferred to a 25 cm² culture flask containing an additional 3.5 ml of medium. For overnight recovery, cells were kept at 37°C and 5% CO₂. On the following day, the numbers of viable and dead cells were determined by trypan blue staining and brightfield cell counting. For the time-course stimulation, 5–7 × 10^5^ viable cells per well were seeded into a 24-well plate. PMA (Phorbol 12-myristate 13-acetate; 5 ng/ml) and ionomycin (500 ng/ml) were added to the culture medium, resulting in a total volume of 500 µl per well. PMA functions as an activator of protein kinase C, while ionomycin acts as a calcium ionophore to stimulate immune cell activation and the production of cell type-specific cytokines (Ai et al. 2013; Baran et al. 2001). Particularly considered to induce T cell activity, the addition of PMA/ionomycin has also been demonstrated to stimulate activation signalling in other PBMC subtypes, including B cells, NK cells, dendritic cells, and monocytes (Mandala et al. 2021; Kaszubowska et al. 2018; Ghamlouch et al. 2014; Meyer and Ireland 1998; Saint-Vis et al. 1998). After addition of the stimulants, cells were incubated at 37°C and 5% CO₂. Time-course samples were collected from separate wells before stimulation (0 h; TP0) and at 2 h (TP2) and 4 h (TP4) after stimulant addition.

Fixation and rehydration of the PBMC samples were performed based on the descriptions by Chen et al. (Chen et al. 2018) and the adapted protocol from 10x Genomics, Inc. (Methanol Fixation of Cells for Single Cell RNA Sequencing; Pleasanton, CA, USA). In brief, cells were pelleted (4°C, 6 min, 1,600 rpm) and washed twice using pre-cooled (4°C) 0.04% BSA-PBS solution (BSA; heat shock fraction, pH 7, ≥98%; Sigma-Aldrich, St. Louis, MO, USA). All washing steps were conducted using wide-bore pipette tips. The cell pellet was resuspended in 200 µl of 0.04% BSA-DPBS and dehydrated by stirring and dropwise addition of 800 µl of ice-cold methanol (−20°C). Methanol-fixed cell samples were immediately placed at −20°C and subsequently transferred to −80°C for storage. Cell samples were kept on ice throughout the fixation procedure. Time-course collection (TP0, TP2, TP4) across all 28 donors of the main cohort resulted in a total of 84 samples. Methanol-fixed samples were stored for a period of 4–48 days before single-cell library preparation.

### Activation validation by flow cytometry

As the dominant cell type amongst PBMCs, successful immune stimulation was confirmed by the representation of major T cell subpopulations, i.e. CD4+ and CD8+ cells, and the detection of CD69 surface expression. CD69 is considered an early marker of the T cell activation process (Lopez-Cabrera et al. 1993; Diener et al. 2023). Flow cytometry analyses were carried out for the initial donors (donors 1–4) as examples. When collecting the time-course samples as described above, cellular aliquots were taken, washed twice with 1% FCS-PBS (1,600 rpm, 6 min), and fixed using a 4% paraformaldehyde solution (Sigma-Aldrich, St. Louis, MO, USA) in PBS. The following day, cells were stained with fluorescence-coupled antibodies against CD4 (mouse anti-human; clone RPA-T4 (RUO); RRID:AB_398593; Becton, Dickinson and Company), CD8 (mouse anti-human; clone RPA-T8 (RUO); RRID:AB_2738101; Becton, Dickinson and Company), and CD69 (mouse anti-human; PE; clone TP1.55.3; cat. no. IM1943U; Beckman Coulter, Brea, CA, USA). Resulting signals were detected and evaluated using a FACSCanto II flow cytometer and BD FACSDiva™ software (v6.1.3; Becton, Dickinson and Company, Franklin Lakes, NJ, USA). Between 10^3^ and 10^4^ cells were analysed per reaction. Reference ranges for CD4+ and CD8+ T cell proportions were assessed with reference to Valiathan et al. (Valiathan et al. 2014).

### Single-cell RNA sequencing

Methanol-fixed cell samples were equilibrated on ice for 5 min and then pelleted (1,600 rpm, 6 min, 4°C). For rehydration, the cell pellet was resuspended in 25 µl of cold saline sodium citrate (SSC) mix consisting of 3× SSC (diluted from 20× concentrate; Sigma-Aldrich), 0.04% BSA (UltraPure, 50 mg/ml; Thermo Fisher Scientific Inc.), 0.2 U/µl RNase Inhibitor (Protector, 40 U/µl; Sigma-Aldrich), 1 mM DL-dithiothreitol solution (BioUltra, 1 M; Sigma-Aldrich), and nuclease-free water (Invitrogen, Thermo Fisher Scientific Inc.). Resulting cell concentrations were determined from brightfield counting of a sample aliquot stained with trypan blue solution (1:4 dilution).

Single-cell libraries were prepared using the 10x Genomics Chromium Next GEM Single Cell 3′ Reagent Kit v3.1 (PN-1000268) on a Chromium Controller (GCG-SR-1), with the Chromium Next GEM Chip G Single Cell Kit (PN-1000120) and Dual Index Kit TT Set A (PN-1000215; 10x Genomics, Inc., Pleasanton, CA, USA), following the manufacturer’s protocol for methanol-fixed cells (Chen et al. 2018; Zheng et al. 2017). For the pilot cohort (donors 1–4; n = 16 samples), 6,000 cells per sample were loaded; for the main cohort (n = 84 samples), 4,500 cells per sample were loaded. Library quality was assessed using the Agilent High Sensitivity DNA Kit on an Agilent 2100 Bioanalyzer (Agilent Technologies, Santa Clara, CA, USA). Libraries were quantified using the Invitrogen Collibri Library Quantification Kit on a QuantStudio 3 Real-Time PCR System (Thermo Fisher Scientific Inc.). Libraries were pooled equimolarly in batches of 16–18 per lane and sequenced on a NovaSeq S4 instrument using paired-end 150 bp sequencing (PE150) by Novogene (UK) Co., Ltd. (Cambridge, UK).

One sample (PD9_TP0) was flagged as suspect, and a replacement library (PD9_TP0_2) was prepared; the original sample was excluded owing to elevated ambient RNA contamination (see Preprocessing and quality control).

Raw reads were aligned to the human reference genome using Cell Ranger (v7.0.0) and the 10x Genomics refdata-gex-GRCh38-2020-A reference package (GRCh38, released July 2020), for both the pilot and the main cohort.

### Preprocessing and quality control

Per-sample count matrices were loaded into R (v4.0.5) using Seurat (v4.1.1; (Hao et al. 2021)). Ambient RNA contamination was estimated and removed using SoupX (v1.5.2; (Young and Behjati 2020)) with automatic estimation of the contamination fraction. Doublets were identified using DoubletFinder (v2.0.3; (McGinnis et al. 2019)) with an expected doublet rate scaled to the number of cells loaded per sample; the optimal pK parameter was selected by maximising the mean-variance normalised bimodality coefficient. Cells were retained if they had between 200 and 2,500 genes detected and fewer than 10% mitochondrial reads (median 2,197 cells per sample, range 654–6,691; per-sample statistics in Supplementary Figure 1).

### Data integration and cell type annotation

Preprocessed samples were merged and normalised using NormalizeData followed by ScaleData in Seurat (log-normalisation). The top 2,000 highly variable features were selected via FindVariableGenes using variance-stabilizing transformation (vst). Principal component analysis (RunPCA, default parameters beyond npcs = 100) was performed on the scaled feature matrix, and batch effects (by sample identity, orig.ident) were corrected using Seurat’s standard anchor-based integration workflow. The first 30 principal components of the integrated embedding were used for nearest-neighbour graph construction (FindNeighbors, dims 1:30), Louvain community detection (FindClusters, resolution 0.8), and UMAP (RunUMAP, dims 1:30).

Cell type labels were assigned by rule-based, cluster-based manual annotation (Supplementary Table 5) using canonical lineage marker genes. The final annotation comprised 23 cell types across six lineages: CD4 T cells, CD8 T cells, NK cells, B cells, monocytes, and MAIT cells. Cells were assigned hierarchically: major lineage first (Step 1, whole integrated object), then lineage-specific re-clustering and rule-based subtyping within each lineage (Steps 2-5; T cells were further split into CD4/CD8 before subtyping). Annotation quality was assessed by marker gene expression (Supplementary Figure 2) and comparison with known surface-marker hierarchies.

The pilot cohort (donors 1–4) used a different approach: cell-type labels were transferred from the main cohort’s annotated object via Seurat label transfer (FindTransferAnchors, LogNormalize, PCA reduction, dims 1:30; TransferData with the fine annotation as the reference label).

### Tem-CD8 subclustering

Cluster separation quality was assessed by silhouette scoring (cluster::silhouette(), mean silhouette width) across all annotated fine-grained cell types. Tem-CD8 cells (pooled across HC, PD, and NC) were re-clustered on their own variance structure: highly variable genes and PCA (20 PCs) were recomputed within the Tem-CD8 subset alone, uncorrected for patient (not a dominant variance source here) and deliberately not corrected for Timepoint, since that would remove exactly the activation-emergent state this analysis looks for. Clustering resolution (0.1) was set empirically to yield 4 stable, well-separated subclusters.

### Differential gene expression analysis

Differential gene expression was analysed using a pseudobulk approach throughout: raw UMI counts were summed per donor per cell type to form pseudobulk profiles, with no minimum-cell-count threshold at this aggregation step. Genes were considered differentially expressed at a Benjamini-Hochberg FDR < 0.05 and |log₂FC| > 0.5 throughout.

### Timepoint-stratified group comparisons (unpaired)

For comparisons between different individuals at a single stimulation timepoint (PD vs. HC at TP0, TP2, and TP4, sex-stratified PD vs. HC comparisons, and PD vs. NC comparisons), pseudobulk profiles were aggregated using muscat (v1.6.x; (Crowell et al. 2020)) and tested with limma-voom (limma v3.48.x; edgeR v3.34.x; R v4.1.x; (Law et al. 2014; Ritchie et al. 2015; Robinson et al. 2010)): model formula ∼ 0 + condition with a pairwise contrast between groups, library sizes normalised by TMM (edgeR::calcNormFactors() default), gene filtering via muscat::pbDS()’s internal equivalent of edgeR::filterByExpr().

### TBX21/RORC co-expression (HC activation section)

The proportion of TBX21+RORC+ double-positive T cells at each timepoint was compared to TP0 by Fisher’s exact test on a pooled, cell-level contingency table, with the odds ratio and p-value reported directly.

### Within-group activation trajectories (paired)

For comparisons of the same donors across two timepoints within a fixed group (HC, PD, NC, and their sex-stratified equivalents; TP0→TP2, TP0→TP4, TP2→TP4), pseudobulk profiles were aggregated per donor × timepoint and analysed with a paired (patient-blocked) limma-voom design, ∼ donor + timepoint; donors missing either timepoint for a given cell type were excluded from that comparison. Gene filtering applies edgeR::filterByExpr() explicitly (default thresholds: minimum count of 10 reads in a worthwhile number of samples, minimum total count 15) on the pseudobulk (donor × timepoint) library.

PD-vs-HC differences in the paired estimate were tested with a difference-of-coefficients test using the moderated SE saved alongside each effect size (limma’s t = logFC/SE): 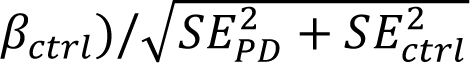, referred to a Welch–Satterthwaite t-distribution, BH-adjusted across genes and cell types.

HC and NC estimates were additionally pooled by fixed-effect inverse-variance-weighted meta-analysis to give a pooled control (n=14) for the same test in place of HC alone (n=5), reported as a power-recovery check rather than independent replication.

Cell types with < 2 complete donor pairs were dropped; those at exactly 2 pairs (below the “reliable” threshold of 3) kept their effect size and SE but their significance score was not calculated.

### Statistical power and minimum detectable effect size

The minimum detectable effect size (MDES) at 80% power and α = 0.05 was computed for a two-sample t-test from the noncentral t-distribution (solved numerically for the effect size at the target power). The pooled SD of the log₂FC effect size (*σ*) was estimated from the median per-gene limma-voom moderated SE for the PD-vs-HC comparisons (TP0/TP2/TP4), back-transformed via 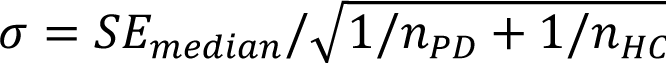.

**Supplementary Figure 11:**
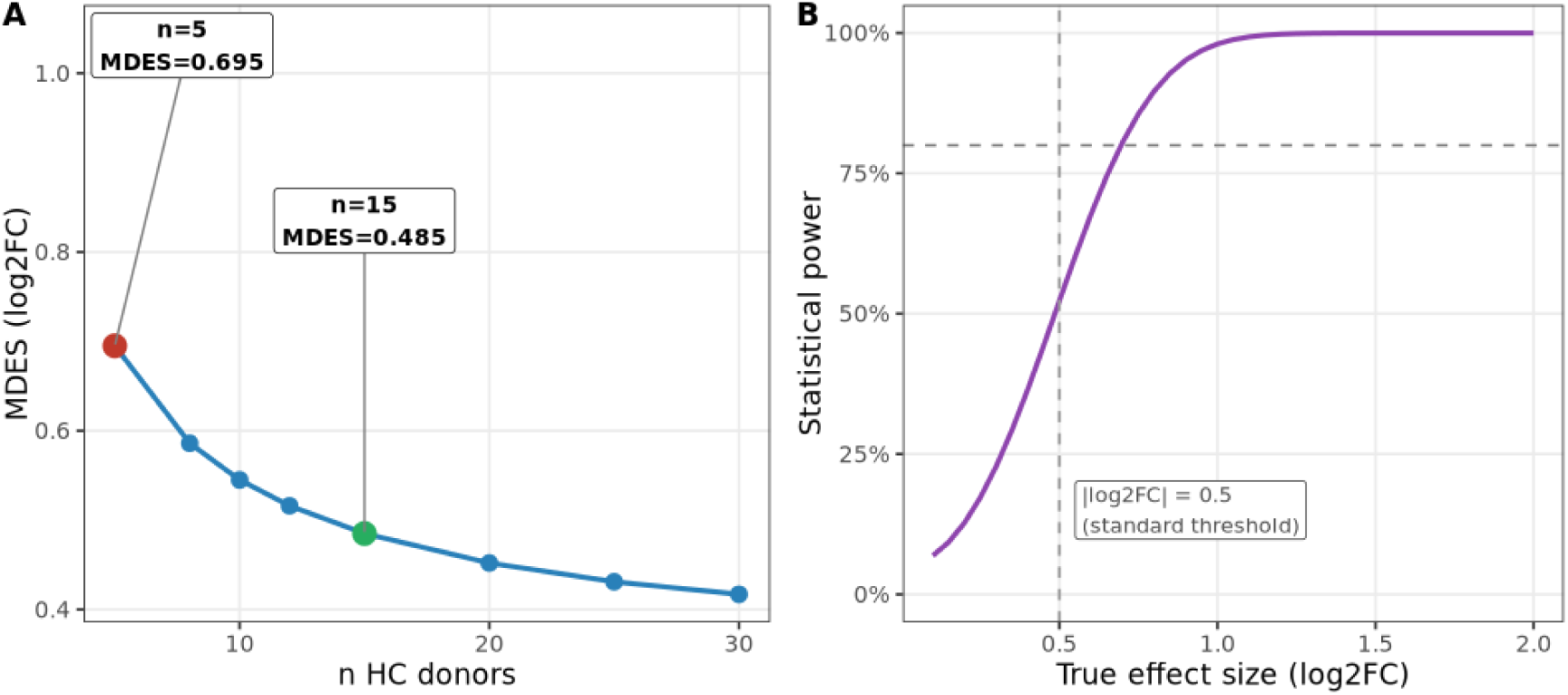
Power analysis for PD vs HC differential expression. (A) Minimum detectable effect size (MDES, log2FC units; 80% power, alpha=0.05) as a function of HC sample size, with PD fixed at n=14. Current n=5 HC and target n=15 HC are marked. (B) Statistical power as a function of true effect size (log2FC) at current n=14 PD / 5 HC. Estimated sigma from the median per-gene limma-voom SE across the PD-vs-HC comparisons, back-transformed to a per-sample SD (Methods).

The unpaired DEG export lacks an SE column, so the median SE was approximated as |log2FC| / qnorm(1 -p/2) from the available log2FC and p-value columns. At current sample sizes (n=14 PD, n=5 HC), the minimum detectable effect size at 80% power is 0.7 log2FC (Supplementary Figure 11 A); the standard |log₂FC| > 0.5 threshold is only 52% powered under these conditions (Supplementary Figure 11 B).

### Cross-study consistency of PD-associated T cell genes

Candidate genes were selected computationally, genome-wide, per lineage (CD4, CD8, NK/NKT): a gene qualifies if it is significant in our own data at one timepoint with at least one other timepoint agreeing in direction, and that direction is also significant in every well-powered external comparator available for that lineage (Bakrac et al. and Wang et al. for CD4; Wang et al. for CD8; Grandke et al., the only dataset profiling NK cells, for NK). Own-data significance is nominal (limma-voom, p < 0.05); external-dataset significance is each dataset’s own within-comparison adjusted p-value (as reported in the original publications).

### Exploratory ligand–receptor bridging to brain tissue

Upregulated ligand genes from our T/NK cell compartments (PD vs. HC at TP0/TP2/TP4, nominal p < 0.05) were matched via directed ligand-receptor pairs from OmniPath (ligrecextra dataset) to receptor genes passing each study’s own reported significance threshold (adjusted p/q < 0.05) in three independent PD brain snRNA-seq studies (Smajic, Martirosyan, and Zhu external datasets). Hits were summarised by ligand × sender cell type × receptor × brain cell type × direction, ranked by the number of confirming brain studies and ligand log₂FC.

### Disease severity correlation (Hoehn-Yahr Stadium)

Per-patient pseudobulk expression was correlated (Pearson) with Hoehn-Yahr Stadium within PD patients, independently per gene, per fine-grained cell type, and per stimulation timepoint. The gene universe tested was restricted to genes already appearing in the FC-and p-value-filtered single-cell Wilcoxon DEG table, not the full transcriptome. A correlation was computed only where at least 5 patients had non-missing expression and Stadium values and both variables had non-zero variance. A gene–cell-type pair was considered part of the consistent Stadium-associated signature if, with n ≥ 10 patients, it showed the same-signed correlation with |r| > 0.5 and nominal p < 0.05 independently at all three timepoints. The same correlation was repeated separately within male and female PD patients for a subset of genes.

### Variance decomposition (PVCA)

Principal variance component analysis (PVCA; (Li et al. 2009)) was performed using the R pvca package (with lme4 and Biobase) on pseudobulk expression aggregated per sample by summing raw counts (Seurat AggregateExpression()), restricted to the top 2,000 most variable genes by variance and log2(count + 1)-transformed (a simple log transform of the pseudobulk sum, not a library-size-normalised or voom-transformed value). A combined-timepoint model (pvcaBatchAssess() threshold 0.6) was performed separately within each disease group (HC, PD, NC) using a full model of the covariates with more than one level and fewer levels than samples in that group (PD: Group_Detail, Timepoint, clinical_type_description, Stadium, sex; HC and NC: Timepoint, sex only).

A separate, per-timepoint PVCA (threshold 0.75; sex, clinical_type_description, Group_Detail, Stadium as covariates) was performed to test the same variance decomposition at each timepoint individually.

### Cross-sex concordance

Cross-sex fold-change concordance was quantified by Pearson correlation of per-gene log₂FC values between sex-stratified DEG results (single-cell Wilcoxon avg_log2FC, chosen for complete cell-type × timepoint coverage independent of pseudobulk replicate count), computed per cell type at each timepoint. Because each sex-stratified stratum contains at most seven donors, we treat Pearson r as the primary descriptive statistic; BH-adjusted p-values (two-sided t-test on Fisher-z transformed r) are reported alongside for context. To confirm that the high male–female concordance observed in Tem-CD8 at TP4 specifically exceeds chance, a permutation null distribution was constructed by shuffling gene labels within one sex’s log₂FC vector 500 times (seed fixed at 42) and recomputing the Pearson correlation; the observed correlation was compared against the 95th percentile of this null distribution.

### Pathway analysis

#### Pre-ranked GSEA (pathway-level activation dynamics)

To identify the top enriched Hallmark pathways per activation phase in HC (Results, “The normal activation programme in healthy PBMCs”), pre-ranked GSEA was performed with GSEApy (gseapy.prerank, v1.1.3; Python v3.10, pandas v1.5.3, numpy v1.23.5) on paired, patient-blocked limma-voom pseudobulk differential expression results, run independently for each cell type × comparison group. Genes were ranked directly by signed avg_log2FC (no additional transformation); genes below |log2FC| = 0.15 were dropped before ranking, and the ranked list was trimmed to the top 800 genes by |log2FC| per group for tractability. Gene sets were the union of three MSigDB-style collections supplied as local .gmt files: GO Biological Process (go_bp), Hallmark (hallmarks), and ImmuneSigDB (immunesig); gene sets with fewer than 10 or more than 2,000 members were excluded. Significance was assessed by permutation testing (250 permutations per group, seed fixed at 42); pathways were considered enriched at FDR (BH-adjusted permutation q-value) < 0.25 (Subramanian et al. 2005).

#### Over-representation analysis (curated gene sets)

For targeted questions operating on a specific, pre-defined gene list rather than a full ranked comparison (the Stadium-severity-correlated gene programme per cell type, the sex-specific PD-baseline gene sets, and the shared/PD-specific/Neurological Control-specific DEG sets at TP4), over-representation of Gene Ontology Biological Process terms was tested using clusterProfiler::enrichGO() against org.Hs.eg.db (SYMBOL keytype; clusterProfiler v4.10.0, org.Hs.eg.db v3.18.0, R v4.3.1; (Wu et al. 2021)) against a comparison-specific background (all genes tested in that analysis), with Benjamini-Hochberg-adjusted q < 0.05. The top five terms by q-value are reported per gene set, terms are additionally required to have Count ≥ 3 (at least 3 of the input genes hitting that term).

### Module scores

Module scores (mean normalised expression of curated gene sets) were computed per cell using the mean of normalised expression values across all detected module genes, then averaged per donor × timepoint × cell type. Every curated gene panel used in this manuscript is listed in Supplementary Table 6.

#### CD8 T cell senescence score is a signed composite, not a plain mean

CDKN1A, CDKN2A, B3GAT1, and KLRG1 increase with T cell senescence, while CD27 and CD28 are lost. To account for this, the score is mean(CDKN1A, CDKN2A, B3GAT1, KLRG1) minus mean(CD27, CD28), with each sub-score independently z-scaled (as in the resolution-failure test below) before differencing, rather than a single unsigned mean across all six genes. CD57 (B3GAT1) discriminates proliferative incapacity more specifically than CD28 loss alone (CD57 ^+^ CD28 ^+^ cells are already non-proliferative) (Brenchley et al. 2003), though it also tracks cumulative antigen exposure/CMV serostatus alongside chronological senescence (Autaa et al. 2025); both are retained as the field’s standard markers.

#### Paired module-score effect sizes (resolution-failure test)

To test whether a group’s activation-induced rise in a module score differs between the early (TP0→TP2) and late (TP2→TP4) timepoint windows, module scores (pseudobulk, z-scaled within cell type) were differenced within donor to give a paired delta per window. Within-group effect size is Cohen’s dz (mean paired delta / SD of the paired delta); between-group effect size is Cohen’s d comparing PD’s deltas to HC’s deltas in that window, 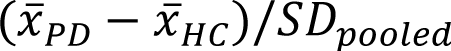 Cell types are ranked by a “resolution-failure” score (|between-group d| in the late window minus the early window) rather than by within-group significance, since HC’s n=5 makes within-group Wilcoxon tests under-powered by construction; effect size therefore drives the ranking, with between-group p-values (two-sample Wilcoxon, BH-adjusted across cell types within each window) reported alongside for reference. Cell types with fewer than 3 donor pairs in either group for a given window are excluded.

### Differential cell type abundance

Cell type composition was quantified per sample (donor × timepoint) as raw cell counts per fine-grained cell type, centred-log-ratio (CLR) transformed (count + 0.5 pseudocount; divided by the per-sample geometric mean across cell types, then log-transformed) to respect the compositional (sum-constrained) nature of cell type proportions.

#### Within-group activation trajectories

For the longitudinal, within-group comparisons (HC, PD, and NC each compared across their own TP0→TP2→TP4 trajectory), the same CLR-transformed composition was modelled per cell type per group with donor included as a fixed blocking effect, CLR(count) ∼ patient + Timepoint, mirroring the paired-samples design adopted for the DEG pipeline earlier. Both timepoint coefficients (TP2 vs. TP0, TP4 vs. TP0) were extracted from the same model fit; BH adjustment was applied once, globally, across all cell types, groups, and both timepoint terms together. Results were computed separately within each sex.

#### Tem-CD8 exhaustion subcluster (cross-sectional, TP4)

The same CLR-transformed compositional framework (pseudocount 0.5, geometric mean across categories) was applied with the Tem-CD8 subclusters as the category system and one row per patient × timepoint, restricted to TP4 and to the exhaustion-associated subcluster. A linear model, CLR(proportion) ∼ Group + age (Group: HC, NC, PD), was fit.

### Cell-cell communication analysis (CellChat)

Cell-cell communication was inferred using CellChat (Jin et al. 2021, 2025) on normalised expression data. Communication was inferred with the human ligand-receptor database (CellChatDB.human). For each comparison, CellChat’s standard pipeline was applied: subsetData(), identifyOverExpressedGenes() (exhaustive search, do.fast = FALSE), identifyOverExpressedInteractions(), computeCommunProb() (type = “triMean”), filterCommunication() (minimum 10 cells per group), computeCommunProbPathway(), and aggregateNet(). PD vs. HC was compared at each timepoint independently (TP0, TP2, TP4), each requiring a minimum of 50 cells per group/timepoint to proceed. Per-comparison CellChat objects were merged (mergeCellChat), and ligand-receptor-level interaction probabilities and per-pathway summed probabilities (used for the “communication probability”/“total probability” metrics reported in the text and figures) were extracted via subsetCommunication(). Ligand-receptor pairs were filtered to CellChat’s own permutation-based significance threshold (p < 0.05).

### External datasets used

All comparator datasets below were obtained as published differential-expression results from each study’s own supplementary materials, not reprocessed from raw data.

#### Cross-study DEG replication (peripheral blood)

| Dataset | Data type / tissue | Celltypes |
| --- | --- | --- |
| Meglaj Bakrač et al.<br>(Meglaj Bakrač et al. 2026) | scRNA-seq, PD + HC PBMC | CD4 T cells |
| Wang et al. (Wang et al. 2021) | scRNA-seq, PD + HC blood | CD4 and CD8 T cells |
| Grandke et al. (Grandke et al. 2025) | scRNA-seq, PD + HC PBMC | Unsorted PBMCs |

#### Exploratory ligand-receptor bridging (PD brain tissue)

| Dataset | Data type / tissue |
| --- | --- |
| Smajić et al. (Smajić et al. 2022) | snRNA-seq, midbrain/substantia nigra |
| Martirosyan et al.<br>(Martirosyan et al. 2024) | snRNA-seq, substantia nigra pars compacta |
| Zhu et al. (Zhu et al. 2024) | snRNA-seq, prefrontal cortex |

### Statistical reporting

Analyses were performed in R v4.0.5 with Seurat v4.1.1 for preprocessing, data integration, and clustering. The GSEA step (gseapy v1.1.3) runs in Python v3.10. Statistical tests are identified in figure legends and described in the relevant Methods subsections above; all tests are two-sided unless a figure legend explicitly states otherwise, and multiple testing correction used the Benjamini-Hochberg procedure unless otherwise stated. Boxplots throughout follow the standard Tukey convention: the box spans the 25th–75th percentile (interquartile range, IQR), the centre line marks the median, and whiskers extend to the most extreme data point within 1.5×IQR of the box; individual data points (patients/donors) are overlaid directly rather than plotted as a separate outlier layer (`outlier.shape = NA`). Shaded ribbons and error bands represent either the 95% confidence interval of a fitted linear model or the mean ± standard error of the mean (SEM = SD/√n) for per-timepoint trajectory summaries; each figure legend specifies which. Correlation-based concordance and replication analyses throughout this manuscript (cross-sex, cross-cohort, cross-disease, and technical-validation checks) use Pearson rather than Spearman correlation, since these analyses ask whether two conditions or cohorts show quantitatively comparable fold-change magnitudes (not merely the same gene ranking). Summary statistics are reported as mean (range) for cohort demographics and median (IQR) for distributional summaries. Sample sizes for each comparison are stated directly in the relevant figure legend; full per-patient sample metadata (group, sex, timepoint, Hoehn-Yahr Stadium) is given in Supplementary Table 1.

## Acknowledgements

Special thanks to Sébastien Dorison and Dr. Jakob Stögbauer (University Hospital of Saarland, Department of Neurology) for their excellent support with clinical matters.

## Author contributions

Study design: C.D., F.G., M.H., N.L., M.U., K.F., A.K. and E.M. Donor recruiting and blood collection: B.B. and A.B.-D. with support from K.-U.D. PBMC isolation: B.B. and C.D. with support from B.W.-R. and T.T. Time-course experiments and cell fixation: C.D. with support from B.B. Antibody staining and FACS analysis: T.T., C.D. and B.W.-R. Library generation and preparations for sequencing analysis: C.D. with support from N.L. Bioinformatics analyses: F.G. Manuscript writing: F.G. and C.D.

## Declaration of generative AI and AI-assisted technologies

During the preparation of this work, the author(s) used Claude (Anthropic) to assist with drafting and editing manuscript text. After using this tool, the author(s) reviewed and edited the content as needed and take full responsibility for the content of the published work.

## References

1. Ai, Wenchao, Haishan Li, Naining Song, Lei Li, and Huiming Chen. 2013. “Optimal Method to Stimulate Cytokine Production and Its Use in Immunotoxicity Assessment.” International Journal of Environmental Research and Public Health 10 (9): 3834–42.

2. Autaa, Gaëlle, Daniil Korenkov, Josine Van Beek, et al. 2025. “Re-Evaluating CD57 as a Marker of t Cell Senescence: Implications for Immune Ageing and Differentiation.” Immunity & Ageing 22 (1): 47.

3. Baek, Jean-Ha, Dejan Mamula, Beata Tingstam, Marcela Pereira, Yachao He, and Per Svenningsson. 2019. “GRP78 Level Is Altered in the Brain, but Not in Plasma or Cerebrospinal Fluid in Parkinson’s Disease Patients.” Frontiers in Neuroscience 13: 697.

4. Baird, Jill K., Dennis Bourdette, Charles K. Meshul, and Joseph F. Quinn. 2019. “The Key Role of t Cells in Parkinson’s Disease Pathogenesis and Therapy.” Parkinsonism & Related Disorders 60: 25–31.

5. Baran, Jarołsaw, Danuta Kowalczyk, Mariola Ożóg, and Marek Zembala. 2001. “Three-Color Flow Cytometry Detection of Intracellular Cytokines in Peripheral Blood Mononuclear Cells: Comparative Analysis of Phorbol Myristate Acetate-Ionomycin and Phytohemagglutinin Stimulation.” Clinical Diagnostic Laboratory Immunology 8 (2): 303–13.

6. Bautista, Juan Miguel, Christyn Jezza Quanico, Paulo Cataniag, Jose Gil Guillermo, and Nobutaka Hattori. 2026. “Detecting the Seeds of Parkinson’s Disease: The Evolution of α–synuclein Seed Amplification Assay as a Clinical Biomarker.” Journal of Neural Transmission, 1–7.

7. Blank, Christian U., W. Nicholas Haining, Werner Held, et al. 2019. “Defining ‘t Cell Exhaustion’.” Nature Reviews Immunology 19 (11): 665–74.

8. Bovenzi, Roberta, Matteo Conti, Clara Simonetta, et al. 2025. “Sex-Specific Immune-Biological Profiles in Parkinson’s Disease.” Journal of Neuroimmunology 403: 578610.

9. Brenchley, Jason M., Nitin J. Karandikar, Michael R. Betts, et al. 2003. “Expression of CD57 Defines Replicative Senescence and Antigen-Induced Apoptotic Death of CD8+ t Cells.” *Blood*, The Journal of the American Society of Hematology 101 (7): 2711–20.

10. Brochard, Vanessa, Béhazine Combadière, Annick Prigent, et al. 2009. “Infiltration of CD4+ Lymphocytes into the Brain Contributes to Neurodegeneration in a Mouse Model of Parkinson Disease.” The Journal of Clinical Investigation 119 (1).

11. Capelle, Christophe M., Séverine Ciré, Fanny Hedin, et al. 2023. “Early-to-Mid Stage Idiopathic Parkinson’s Disease Shows Enhanced Cytotoxicity and Differentiation in CD8 t-Cells in Females.” Nature Communications 14 (1): 7461.

12. Cerri, Silvia, Liudmila Mus, and Fabio Blandini. 2019. “Parkinson’s Disease in Women and Men: What’s the Difference?” Journal of Parkinson’s Disease 9 (3): 501–15.

13. Chamorro, Ángel, Andreas Meisel, Anna M. Planas, Xabier Urra, Diederik Van De Beek, and Roland Veltkamp. 2012. “The Immunology of Acute Stroke.” Nature Reviews Neurology 8 (7): 401–10.

14. Chen, Jinguo, Foo Cheung, Rongye Shi, Huizhi Zhou, Wenrui Lu, and C. H. I. Consortium Candia Julián julian. candia@ nih. gov Kotliarov Yuri yuri. kotliarov@ nih. gov Stagliano Katie R. staglianoke@ mail. nih. gov Tsang John S. john. tsang@ nih. gov. 2018. “PBMC Fixation and Processing for Chromium Single-Cell RNA Sequencing.” Journal of Translational Medicine 16 (1): 198.

15. Chen, Joyce, Isaac F. López-Moyado, Hyungseok Seo, et al. 2019. “NR4A Transcription Factors Limit CAR t Cell Function in Solid Tumours.” Nature 567 (7749): 530–34.

16. Collins, Sam, Michael A. Lutz, Paul E. Zarek, Robert A. Anders, Gilbert J. Kersh, and Jonathan D. Powell. 2008. “Opposing Regulation of t Cell Function by Egr–1/NAB2 and Egr–2/Egr–3.” European Journal of Immunology 38 (2): 528–36.

17. Contaldi, Elena, Luca Magistrelli, Anna Vera Milner, Marco Cosentino, Franca Marino, and Cristoforo Comi. 2021. “Expression of Transcription Factors in CD4+ t Cells as Potential Biomarkers of Motor Complications in Parkinson’s Disease.” Journal of Parkinson’s Disease 11 (2): 507–14.

18. Crabtree, Gerald R. 1989. “Contingent Genetic Regulatory Events in t Lymphocyte Activation.” Science 243 (4889): 355–61.

19. Crowell, Helena L., Charlotte Soneson, Pierre-Luc Germain, et al. 2020. “Muscat Detects Subpopulation-Specific State Transitions from Multi-Sample Multi-Condition Single-Cell Transcriptomics Data.” Nature Communications 11 (1): 6077.

20. Davidi, Dana, Meir Schechter, Suaad Abd Elhadi, Adar Matatov, Lubov Nathanson, and Ronit Sharon. 2020. “Α-Synuclein Translocates to the Nucleus to Activate Retinoic-Acid-Dependent Gene Transcription.” IScience 23 (3).

21. Diener, Caroline, Martin Hart, Tim Kehl, et al. 2023. “Time-Resolved RNA Signatures of CD4+ t Cells in Parkinson’s Disease.” Cell Death Discovery 9 (1): 18.

22. Dorsey, E. Ray, Alexis Elbaz, Emma Nichols, et al. 2018. “Global, Regional, and National Burden of Parkinson’s Disease, 1990-2016: A Systematic Analysis for the Global Burden of Disease Study 2016.” The Lancet Neurology 17 (11): 939–53.

23. Dorsey, E. Ray, Todd Sherer, Michael S. Okun, and Bastiaan R. Bloem. 2018. “The Emerging Evidence of the Parkinson Pandemic.” Journal of Parkinson’s Disease 8 (s1): S3–8.

24. Enogieru, Adaze Bijou, Sylvester Ifeanyi Omoruyi, Donavon Charles Hiss, and Okobi Eko Ekpo. 2019. “GRP78/BIP/HSPA5 as a Therapeutic Target in Models of Parkinson’s Disease: A Mini Review.” Advances in Pharmacological and Pharmaceutical Sciences 2019 (1): 2706783.

25. Feske, Stefan. 2007. “Calcium Signalling in Lymphocyte Activation and Disease.” Nature Reviews Immunology 7 (9): 690–702.

26. Fragiadakis, Gabriela K., Zachary B. Bjornson-Hooper, Deepthi Madhireddy, et al. 2022. “Variation of Immune Cell Responses in Humans Reveals Sex-Specific Coordinated Signaling Across Cell Types.” Frontiers in Immunology 13: 867016.

27. Franceschi, Claudio, Silvana Valensin, Francesco Fagnoni, Cristiana Barbi, and Massimiliano Bonafè. 1999. “Biomarkers of Immunosenescence Within an Evolutionary Perspective: The Challenge of Heterogeneity and the Role of Antigenic Load.” Experimental Gerontology 34 (8): 911– 21.

28. Galiano-Landeira, Jordi, Albert Torra, Miquel Vila, and Jordi Bové. 2020. “CD8 t Cell Nigral Infiltration Precedes Synucleinopathy in Early Stages of Parkinson’s Disease.” Brain 143 (12): 3717– 33.

29. Gate, David, Naresha Saligrama, Olivia Leventhal, et al. 2020. “Clonally Expanded CD8 t Cells Patrol the Cerebrospinal Fluid in Alzheimer’s Disease.” Nature 577 (7790): 399–404. 10.1038/s41586-019-1895-7.

30. Ghamlouch, Hussein, Hakim Ouled–Haddou, Aude Guyart, et al. 2014. “Phorbol Myristate Acetate, but Not CD40L, Induces the Differentiation of CLL b Cells into Ab–secreting Cells.” Immunology and Cell Biology 92 (7): 591–604.

31. Goldeck, David, Claudia Schulte, Marcia Cristina Teixeira dos Santos, et al. 2022. “Higher Frequencies of t-Cells Expressing NK-Cell Markers and Chemokine Receptors in Parkinson’s Disease.” Journal of Ageing and Longevity 3 (1): 1–10.

32. Grandke, Friederike, Tobias Fehlmann, Fabian Kern, et al. 2025. “A Single-Cell Atlas to Map Sex-Specific Gene-Expression Changes in Blood Upon Neurodegeneration.” Nature Communications 16 (1): 1965. 10.1038/s41467-025-56833-7.

33. Grozdanov, Veselin, Corinna Bliederhaeuser, Wolfgang P. Ruf, et al. 2014. “Inflammatory Dysregulation of Blood Monocytes in Parkinson’s Disease Patients.” Acta Neuropathologica 128 (5): 651–63.

34. Ham, Tjakko J. van, Mats A. Holmberg, Annemieke T. van der Goot, et al. 2010. “Identification of MOAG-4/SERF as a Regulator of Age-Related Proteotoxicity.” Cell 142 (4): 601–12.

35. Hao, Yuhan, Stephanie Hao, Erica Andersen-Nissen, et al. 2021. “Integrated Analysis of Multimodal Single-Cell Data.” Cell 184 (13): 3573–87.

36. Henson, Sian M., and Arne N. Akbar. 2009. “KLRG1–More Than a Marker for t Cell Senescence.” Age 31 (4): 285–91.

37. Hong, Yanggang, Jingxuan Zhou, Yirong Wang, et al. 2025. “Peripheral Immune Cell-Specific Genes in Parkinson’s Disease Uncovered by Multi-Omics with Therapeutic Implications.” Npj Parkinson’s Disease 11 (1): 302.

38. Isakov, Noah, and Amnon Altman. 2002. “Protein Kinase Cθ in t Cell Activation.” Annual Review of Immunology 20 (1): 761–94.

39. Jin, Suoqin, Christian F. Guerrero-Juarez, Lihua Zhang, et al. 2021. “Inference and Analysis of Cell-Cell Communication Using CellChat.” Nature Communications 12 (1): 1088.

40. Jin, Suoqin, Maksim V. Plikus, and Qing Nie. 2025. “CellChat for Systematic Analysis of Cell–Cell Communication from Single-Cell Transcriptomics.” Nature Protocols 20 (1): 180–219.

41. Kared, Hassen, Serena Martelli, Tze Pin Ng, Sylvia L. F. Pender, and Anis Larbi. 2016. “CD57 in Human Natural Killer Cells and t-Lymphocytes.” Cancer Immunology, Immunotherapy 65 (4): 441– 52.

42. Kaszubowska, Lucyna, Jerzy Foerster, Daria Schetz, and Zbigniew Kmieć. 2018. “CD56bright Cells Respond to Stimulation Until Very Advanced Age Revealing Increased Expression of Cellular Protective Proteins SIRT1, HSP70 and SOD2.” Immunity & Ageing 15 (1): 31.

43. Khan, Omar, Josephine R. Giles, Sierra McDonald, et al. 2019. “TOX Transcriptionally and Epigenetically Programs CD8+ t Cell Exhaustion.” Nature 571 (7764): 211–18. 10.1038/s41586-019-1325-x.

44. Kordower, Jeffrey H., C. Warren Olanow, Hemraj B. Dodiya, et al. 2013. “Disease Duration and the Integrity of the Nigrostriatal System in Parkinson’s Disease.” Brain 136 (8): 2419–31. 10.1093/brain/awt192.

45. Kouli, Antonina, Melanie Jensen, Vanesa Papastavrou, et al. 2021. “T Lymphocyte Senescence Is Attenuated in Parkinson’s Disease.” Journal of Neuroinflammation 18 (1): 228.

46. Kumagai, Tomoaki, Hiroshi Nagayama, Tomohiro Ota, Yasuhiro Nishiyama, Masahiro Mishina, and Masayuki Ueda. 2014. “Sex Differences in the Pharmacokinetics of Levodopa in Elderly Patients with Parkinson Disease.” Clinical Neuropharmacology 37 (6): 173–76.

47. Lang, Benjamin J., Martin E. Guerrero, Thomas L. Prince, Yuka Okusha, Cristina Bonorino, and Stuart K. Calderwood. 2021. “The Functions and Regulation of Heat Shock Proteins; Key Orchestrators of Proteostasis and the Heat Shock Response.” Archives of Toxicology 95 (6): 1943– 70.

48. Law, Charity W., Yunshun Chen, Wei Shi, and Gordon K. Smyth. 2014. “Voom: Precision Weights Unlock Linear Model Analysis Tools for RNA-Seq Read Counts.” Genome Biology 15 (2): R29. 10.1186/gb-2014-15-2-r29.

49. Le, Weidong, Tianhong Pan, Maosheng Huang, et al. 2008. “Decreased NURR1 Gene Expression in Patients with Parkinson’s Disease.” Journal of the Neurological Sciences 273 (1-2): 29–33.

50. Li, Jianying, Pierre R. Bushel, Tzu-Ming Chu, and Russell D. Wolfinger. 2009. “Principal Variance Components Analysis: Estimating Batch Effects in Microarray Gene Expression Data.” Batch Effects and Noise in Microarray Experiments: Sources and Solutions, 141–54.

51. Li, Yi, Ning Chen, Chao Wu, et al. 2020. “Galectin-1 Attenuates Neurodegeneration in Parkinson’s Disease Model by Modulating Microglial MAPK/IκB/NFκB Axis Through Its Carbohydrate-Recognition Domain.” Brain, Behavior, and Immunity 83: 214–25.

52. Lindestam Arlehamn, Cecilia S., Rekha Dhanwani, John Pham, et al. 2020. “Α-Synuclein-Specific t Cell Reactivity Is Associated with Preclinical and Early Parkinson’s Disease.” Nature Communications 11 (1): 1875.

53. Liu, H., L. Wei, Q. Tao, et al. 2012. “Decreased NURR1 and PITX3 Gene Expression in Chinese Patients with Parkinson’s Disease.” European Journal of Neurology 19 (6): 870–75. 10.1111/j.1468-1331.2011.03644.x.

54. Liu, Xindong, Yun Wang, Huiping Lu, et al. 2019. “Genome-Wide Analysis Identifies NR4A1 as a Key Mediator of t Cell Dysfunction.” Nature 567: 525–29. 10.1038/s41586-019-0979-8.

55. Liu, Yan, Hanna K. Sanoff, Hyunsoon Cho, et al. 2009. “Expression of p16INK4a in Peripheral Blood t–cells Is a Biomarker of Human Aging.” Aging Cell 8 (4): 439–48.

56. Lopez-Cabrera, M., A. G. Santis, E. Fernandez-Ruiz, et al. 1993. “Molecular Cloning, Expression, and Chromosomal Localization of the Human Earliest Lymphocyte Activation Antigen AIM/CD69, a New Member of the c-Type Animal Lectin Superfamily of Signal-Transmitting Receptors.” The Journal of Experimental Medicine 178 (2): 537–47.

57. López-Cerdán, Adolfo, Zoraida Andreu, Marta R. Hidalgo, et al. 2022. “Unveiling Sex-Based Differences in Parkinson’s Disease: A Comprehensive Meta-Analysis of Transcriptomic Studies.” Biology of Sex Differences 13 (1): 68.

58. Luo, Yuanrong, Lichun Qiao, Miaoqian Li, Xinyue Wen, Wenbin Zhang, and Xianwen Li. 2025. “Global, Regional, National Epidemiology and Trends of Parkinson’s Disease from 1990 to 2021: Findings from the Global Burden of Disease Study 2021.” Frontiers in Aging Neuroscience 16: 1498756. 10.3389/fnagi.2024.1498756.

59. Macian, Fernando. 2005. “NFAT Proteins: Key Regulators of t-Cell Development and Function.” Nature Reviews Immunology 5 (6): 472–84.

60. Mandala, Wilson, Visopo Harawa, Alinane Munyenyembe, Monica Soko, and Herbert Longwe. 2021. “Optimization of Stimulation and Staining Conditions for Intracellular Cytokine Staining (ICS) for Determination of Cytokine-Producing t Cells and Monocytes.” Current Research in Immunology 2: 184–93.

61. Mark, Julian R., Ann M. Titus, Hannah A. Staley, et al. 2025. “Peripheral Immune Cell Response to Stimulation Stratifies Parkinson’s Disease Progression from Prodromal to Clinical Stages.” Communications Biology 8 (1): 716.

62. Marques, Tainá M., Anouke van Rumund, Ilona B. Bruinsma, et al. 2019. “Cerebrospinal Fluid Galectin-1 Levels Discriminate Patients with Parkinsonism from Controls.” Molecular Neurobiology 56 (7): 5067–74.

63. Martirosyan, Araks, Rizwan Ansari, Francisco Pestana, et al. 2024. “Unravelling Cell Type-Specific Responses to Parkinson’s Disease at Single Cell Resolution.” Molecular Neurodegeneration 19 (1): 1–24.

64. McGinnis, Christopher S., Lyndsay M. Murrow, and Zev J. Gartner. 2019. “DoubletFinder: Doublet Detection in Single-Cell RNA Sequencing Data Using Artificial Nearest Neighbors.” Cell Systems 8 (4): 329–37.

65. Meglaj Bakrač, Sarah, Katarina Mandić, Lidija Cvetko Krajinović, et al. 2026. “Single-Cell Analysis of the Peripheral Immune Landscape in Parkinson’s Disease: Insights into Dendritic Cell and CD4+ t-Cell Transcriptomics.” Npj Parkinson’s Disease 12 (1): 73. 10.1038/s41531-026-01283-1.

66. Meller, Nahum, Amnon Altman, and Noah Isakov. 1998. “New Perspectives on PKCθ, a Member of the Novel Subfamily of Protein Kinase c.” Stem Cells 16 (3): 178–92.

67. Meyer, Kerstin B., and John Ireland. 1998. “PMA/Ionomycin Induces Igκ 3′ Enhancer Activity Which Is in Part Mediated by a Unique NFAT Transcription Complex.” European Journal of Immunology 28 (5): 1467–80.

68. Moquin-Beaudry, Gael, Lovatiana Andriamboavonjy, Sebastien Audet, et al. 2025. “Mapping the Peripheral Immune Landscape of Parkinson’s Disease Patients with Single-Cell Sequencing.” Brain 148 (8): 2847–60. 10.1093/brain/awaf066.

69. Nalls, Mike A., Cornelis Blauwendraat, Costanza L. Vallerga, et al. 2019. “Identification of Novel Risk Loci, Causal Insights, and Heritable Risk for Parkinson’s Disease: A Meta-Analysis of Genome-Wide Association Studies.” The Lancet Neurology 18 (12): 1091–102.

70. Northrop, John K., Rajan M. Thomas, Andrew D. Wells, and Hao Shen. 2006. “Epigenetic Remodeling of the IL-2 and IFN-γ Loci in Memory CD8 t Cells Is Influenced by CD4 t Cells.” The Journal of Immunology 177 (2): 1062–69.

71. Odagiu, Livia, Julia May, Salix Boulet, Troy A. Baldwin, and Nathalie Labrecque. 2021. “Role of the Orphan Nuclear Receptor NR4A Family in t-Cell Biology.” Frontiers in Endocrinology 11: 624122.

72. Pahl, Heike L. 1999. “Activators and Target Genes of Rel/NF-κB Transcription Factors.” Oncogene 18 (49): 6853–66.

73. Parish, Stanley T., Jennifer E. Wu, and Rita B. Effros. 2010. “Sustained CD28 Expression Delays Multiple Features of Replicative Senescence in Human CD8 t Lymphocytes.” Journal of Clinical Immunology 30 (6): 798–805.

74. Pépin, Élise, Tim Jalinier, Guillaume L. Lemieux, Guy Massicotte, and Michel Cyr. 2020. “Sphingosine-1-Phosphate Receptors Modulators Decrease Signs of Neuroinflammation and Prevent Parkinson’s Disease Symptoms in the 1-Methyl-4-Phenyl-1, 2, 3, 6-Tetrahydropyridine Mouse Model.” Frontiers in Pharmacology 11: 77.

75. Pike, Steven C., Matthew Havrda, Francesca Gilli, Ze Zhang, and Lucas A. Salas. 2024. “Immunological Shifts During Early-Stage Parkinson’s Disease Identified with DNA Methylation Data on Longitudinally Collected Blood Samples.” Npj Parkinson’s Disease 10 (1): 21.

76. Pras, Anita, Bert Houben, Francesco A. Aprile, et al. 2021. “The Cellular Modifier MOAG–4/SERF Drives Amyloid Formation Through Charge Complementation.” The EMBO Journal 40 (21): EMBJ2020107568.

77. Ramello, Maria C., Nicolás G. Núñez, Jimena Tosello Boari, et al. 2021. “Polyfunctional KLRG-1+ CD57+ Senescent CD4+ t Cells Infiltrate Tumors and Are Expanded in Peripheral Blood from Breast Cancer Patients.” Frontiers in Immunology 12: 713132.

78. Rao, Anjana, Chun Luo, and Patrick G. Hogan. 1997. “Transcription Factors of the NFAT Family: Regulation and Function.” Annual Review of Immunology 15 (1): 707–47.

79. Recinto, Sherilyn Junelle, Janna E. Jernigan Posey, Nadejda Lefter, et al. 2026. “Profiling Peripheral Immune Cells in Parkinson’s Disease: A Scoping Review.” bioRxiv, 2026–02.

80. Ritchie, Matthew E., Belinda Phipson, D. I. Wu, et al. 2015. “Limma Powers Differential Expression Analyses for RNA-Sequencing and Microarray Studies.” Nucleic Acids Research 43 (7): e47–47.

81. Robinson, Mark D., Davis J. McCarthy, and Gordon K. Smyth. 2010. “edgeR: A Bioconductor Package for Differential Expression Analysis of Digital Gene Expression Data.” Bioinformatics 26 (1): 139–40.

82. Rocha, Natália Pessoa, Antônio Lúcio Teixeira, Paula Luciana Scalzo, et al. 2014. “Plasma Levels of Soluble Tumor Necrosis Factor Receptors Are Associated with Cognitive Performance in Parkinson’s Disease.” Movement Disorders 29 (4): 527–31.

83. Saint-Vis, Blandine de, Isabelle Fugier-Vivier, Catherine Massacrier, et al. 1998. “The Cytokine Profile Expressed by Human Dendritic Cells Is Dependent on Cell Subtype and Mode of Activation.” The Journal of Immunology 160 (4): 1666–76.

84. Salemi, Michele, Filomena Cosentino, Giuseppe Lanza, et al. 2021. “mRNA Expression Profiling of Mitochondrial Subunits in Subjects with Parkinson’s Disease.” Archives of Medical Science: AMS 19 (3): 678.

85. Shaulian, Eitan, and Michael Karin. 2002. “AP-1 as a Regulator of Cell Life and Death.” Nature Cell Biology 4 (5): E131–36.

86. Smajić, Semra, Cesar A. Prada-Medina, Zied Landoulsi, et al. 2022. “Single-Cell Sequencing of Human Midbrain Reveals Glial Activation and a Parkinson-Specific Neuronal State.” Brain 145 (3): 964–78.

87. Subramanian, Aravind, Pablo Tamayo, Vamsi K. Mootha, et al. 2005. “Gene Set Enrichment Analysis: A Knowledge-Based Approach for Interpreting Genome-Wide Expression Profiles.” Proceedings of the National Academy of Sciences 102 (43): 15545–50.

88. Sulzer, David, Roy N. Alcalay, Francesca Garretti, et al. 2017. “T Cells from Patients with Parkinson’s Disease Recognize α-Synuclein Peptides.” Nature 546 (7660): 656–61.

89. Sun, Fei, Tian-Tian Yue, Chun-Liang Yang, et al. 2021. “The MAPK Dual Specific Phosphatase (DUSP) Proteins: A Versatile Wrestler in t Cell Functionality.” International Immunopharmacology 98: 107906.

90. Tansey, Malú Gámez, Rebecca L. Wallings, Madelyn C. Houser, Mary K. Herrick, Cody E. Keating, and Valerie Joers. 2022. “Inflammation and Immune Dysfunction in Parkinson Disease.” Nature Reviews Immunology 22 (11): 657–73.

91. Tranchevent, Léon-Charles, Rashi Halder, and Enrico Glaab. 2023. “Systems Level Analysis of Sex-Dependent Gene Expression Changes in Parkinson’s Disease.” Npj Parkinson’s Disease 9 (1): 8.

92. Truneh, Alemseged, Françoise Albert, Pierre Golstein, and Anne-Marie Schmitt-Verhulst. 1985. “Early Steps of Lymphocyte Activation Bypassed by Synergy Between Calcium Ionophores and Phorbol Ester.” Nature 313 (6000): 318–20.

93. Tumpa, Jarika Jahan, Qurat Ul Ain Hayder, Nowshin Sharmily Maisa, and Md Nazmul Islam. 2025. “Peripheral-Central Immune Interactions in Parkinson’s Disease: Insights into Innate and Adaptive Immunity.” Frontiers in Molecular Neuroscience 18: 1682006.

94. Uehara, Takashi, Tomohiro Nakamura, Dongdong Yao, et al. 2006. “S-Nitrosylated Protein-Disulphide Isomerase Links Protein Misfolding to Neurodegeneration.” Nature 441 (7092): 513–17.

95. Valiathan, Ranjini, Khaled Deeb, Marc Diamante, Margarita Ashman, Naresh Sachdeva, and Deshratn Asthana. 2014. “Reference Ranges of Lymphocyte Subsets in Healthy Adults and Adolescents with Special Mention of t Cell Maturation Subsets in Adults of South Florida.” Immunobiology 219 (7): 487–96.

96. Vallabhapurapu, Sivakumar, and Michael Karin. 2009. “Regulation and Function of NF-κB Transcription Factors in the Immune System.” Annual Review of Immunology 27 (1): 693–733.

97. Wang, Pingping, Lifen Yao, Meng Luo, et al. 2021. “Single-Cell Transcriptome and TCR Profiling Reveal Activated and Expanded t Cell Populations in Parkinson’s Disease.” Cell Discovery 7 (1): 52.

98. Wang, Quan, and Qun Xue. 2023. “Bioinformatics Analysis of Potential Common Pathogenic Mechanism for Carotid Atherosclerosis and Parkinson’s Disease.” Frontiers in Aging Neuroscience 15: 1202952.

99. Wherry, E. John, Sang-Jun Ha, Susan M. Kaech, et al. 2007. “Molecular Signature of CD8+ t Cell Exhaustion During Chronic Viral Infection.” Immunity 27 (4): 670–84.

100. Wherry, E. John, and Makoto Kurachi. 2015. “Molecular and Cellular Insights into t Cell Exhaustion.” Nature Reviews Immunology 15 (8): 486–99.

101. Williams-Gray, C. H., R. S. Wijeyekoon, K. M. Scott, S. Hayat, R. A. Barker, and J. L. Jones. 2018. “Abnormalities of Age-Related t Cell Senescence in Parkinson’s Disease.” Journal of Neuroinflammation 15 (1): 166.

102. Wu, Tianzhi, Erqiang Hu, Shuangbin Xu, et al. 2021. “clusterProfiler 4.0: A Universal Enrichment Tool for Interpreting Omics Data.” The Innovation 2 (3).

103. Xiong, Liu-Lin, Ruo-Lan Du, Rui-Ze Niu, et al. 2024. “Single-Cell RNA Sequencing Reveals Peripheral Immunological Features in Parkinson’s Disease.” Npj Parkinson’s Disease 10 (1): 185.

104. Xu, Weili, and Anis Larbi. 2017. “Markers of t Cell Senescence in Humans.” International Journal of Molecular Sciences 18 (8): 1742.

105. Yan, Mingmin, Lanxia Meng, Lijun Dai, et al. 2020. “Cofilin 1 Promotes the Aggregation and Cell-to-Cell Transmission of α-Synuclein in Parkinson’s Disease.” Biochemical and Biophysical Research Communications 529 (4): 1053–60.

106. Yan, Mingmin, Min Xiong, Lijun Dai, et al. 2022. “Cofilin 1 Promotes the Pathogenicity and Transmission of Pathological α-Synuclein in Mouse Models of Parkinson’s Disease.” Npj Parkinson’s Disease 8 (1): 1.

107. Young, Matthew D., and Sam Behjati. 2020. “SoupX Removes Ambient RNA Contamination from Droplet-Based Single-Cell RNA Sequencing Data.” Gigascience 9 (12): giaa151.

108. Zheng, Grace X. Y., Jessica M. Terry, Phillip Belgrader, et al. 2017. “Massively Parallel Digital Transcriptional Profiling of Single Cells.” Nature Communications 8 (1): 14049.

109. Zhu, Biqing, Jae-Min Park, Sarah R. Coffey, et al. 2024. “Single-Cell Transcriptomic and Proteomic Analysis of Parkinson’s Disease Brains.” Science Translational Medicine 16 (771): eabo1997.

110. Zhu, Zhu, Yu–Wen Wang, Ding–Hao Ge, et al. 2017. “Downregulation of DEC 1 Contributes to the Neurotoxicity Induced by MPP+ by Suppressing PI 3K/Akt/GSK 3β Pathway.” CNS Neuroscience & Therapeutics 23 (9): 736–47.

